# Do eyes say it all? Assessing the utility of physiological signals for predicting systemic cognitive states

**DOI:** 10.1101/2024.02.29.582708

**Authors:** Ayca Aygun, Thuan Nguyen, Matthias Scheutz

## Abstract

Robust estimation of systemic human cognitive states is critical for many applications, from simply detecting inefficiencies in human task performance to adapting the behaviors of artificial agents to improve team performance in mixed-initiative human-machine teams. Here we use comprehensive analyses of a multi-modal dataset from a multi-tasking driving experiment to provide systematic evidence that percentage change in pupil size (PCPS) in human eye gaze is the most reliable biomarker for assessing three human cognitive states relevant to task performance: workload, sense of urgency, and mind wandering. Specifically, we performed comprehensive statistical tests that establish the superior performance of PCPS compared other physiological signals like electroencephalogram (EEG), functional near-infrared spectroscopy (fNIRS), respiration, arterial blood pressure (ABP), and skin conductance. Based on the eye gaze data, we also characterized the relation between workload and sense of urgency, observing that consecutive occurrences of higher sense of urgency tended to increase overall workload. We also trained five state-of-the-art machine learning models on the data to determine their potential for predicting the three systemic cognitive states across subjects, showing that four of them had similar highest accuracy in cognitive state classification based on PCPS (with one, random forest, showing inferior performance). In combination with the statistical analyses, the comparative modeling results demonstrate that PCPS, in contrast to prior findings in the literature, represents the optimal method for estimating the three performance-relevant cognitive states.

## Introduction

A reliable determination of human systemic cognitive states, including *cognitive workload*, *sense of urgency*, and *mind wandering*, would be helpful in many task-based settings, and even critical in some cases (*e.g.*, detecting mentally overloaded humans in safety-critical contexts who will be more prone to errors and oversight due to being overloaded, or detecting mind wandering in situations where human observers might miss important events due to attentional drifts). Being able to assess human systemic cognitive states can also be of great utility for artificial agents operating in human-agent teams in that it would allow artificial agents to adapt their behavior and the teams’ task allocation to achieve a better load balance and increase the effectiveness of the whole team (rather than acting in ways that might, for example, increase human cognitive workload [11]). Similarly, it would be important in some contexts to track mind wandering, defined as the detachment of human attention from the current task, to prevent the decline in situational awareness of human performers [12]. Moreover, information about a person’s sense of urgency could be used as a social signal, prompting them to pay attention to social cues in a human-machine interaction setting and help artificial agents to better understand the needs of their human teammates [13].

In this paper, we use a comprehensive dataset collected from an interactive multi-modal multi-task driving experiment [15] to investigate the extent to which the above three systemic human cognitive states (cognitive workload, sense of urgency, and mind wandering) could be inferred from various physiological signals including eye gaze, EEG, fNIRS, skin conductance, and respiration rate. Based on the analyses, we then aimed to develop general computational models that, when trained on a small set of human physiological data, would be able to detect and predict those states in any person.

The two main contributions of this paper are thus:

- a comprehensive investigation of the utilitity of various physiological signals including pupillometry, EEG, fNIRS, skin conductance, and respiration for determining **(1)** workload, **(2)** sense of urgency, and **(3)** mind wandering through various statistical analyzes, which shows the superiority of using “percentage change in pupil size” (PCPS) over all other physiological biomarkers.
- an exploration of five state-of-the-art machine learning (ML) models trained on the data (*k*-Nearest Neighbor (k-NN), Naive-Bayes (NB), Random Forest (RF), Support Vector Machines (SVM), and Multiple Layer Perceptron (MLP)) to evaluate the effectiveness of multiple physiological signal types for detecting and predicting different levels of the three cognitive state levels, showing again that the best prediction results can be obtained with the PCPS.

The remainder of this paper is structured as follows. We start with our motivation for considering the above three human systemic cognitive states as well as a brief description of the data we used for our analysis and modeling efforts, followed by a review of related work on using different sensory modalities to detect them. We then provide details on the employed data and how it was used, including information on data selection and preprocessing for each of the sensory modalities. We then present the statistical analyses together with the performance results of different machine learning models, followed by a discussion of the implications for the broader human-machine teaming context, but also the limitations and the future work. Finally, we conclude with a summary of our accomplishments, highlighting again the utility of collecting and processing eye gaze for assessing human performance.

## Motivation

Human *systemic cognitive states* like workload, sense of urgency, but also mind wandering, can modulate and impact human task performance (e.g., [10]), be it in individual tasks or as part of mixed initiative teams (e.g., [3]). While there is independent foundational interest in computational models for detecting and predicting the presence of performance-relevant cognitive states from a psychological perspective, the important practical application of such models would be to use them for improving human performance, for example, by reassigning tasks based on workload, or redirecting human attention to the task processes during periods of mind wandering. We are particularly interested in performance-relevant systemic states and their possible interactions as they typically do not occur in isolation but can influence each other (see Figure 1), focussing our investgiation on workload, sense of urgency, and mind wandering.

**Fig 1.**
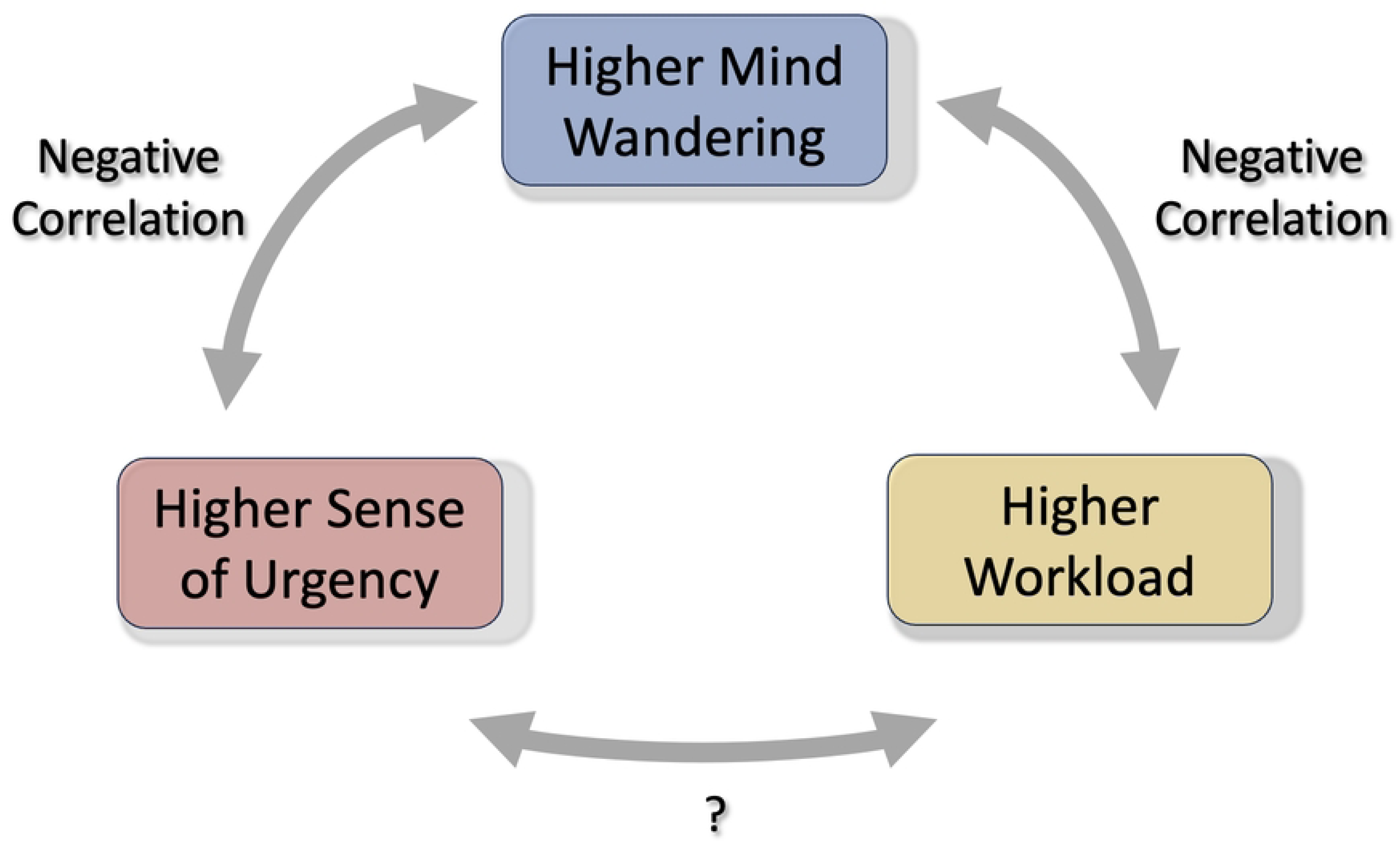
The relationship between Different Cognitive State Pairs. An illustration of the hypothesized relationships among pairs of three systemic cognitive states: workload, sense of urgency, and mind wandering.

### Workload

has been described as a human-centered concept which arises from the interactions between task requirements, the environmental conditions under which the task is performed, as well as the performer’s abilities, attitudes, and awareness [18]. Workload has also been associated with the cognitive demands to meet task requirements [19]. In particular, workload is evaluated as a multidimensional subjective experience that reflects the cost incurred by an operator in achieving a given level of performance, emerging from the interaction between task demands, environmental conditions, and the operator’s skills, behaviors, and perceptions.

### Sense of urgency

has been defined as the perception that a given task must be accomplished quickly, usually within a certain period of time, which affects decision-making mechanisms in humans. For example, Yau *et al.* found that it is possible to identify EEG signals which are related to the sense of urgency [20]. Another study examined the impact of warning signals, including auditory, vibrotactile, and multimodal, in assessing sense of urgency in different cognitive workload conditions [21].

### Mind wandering

has been defined as a particular case of spontaneous thought, a cognitive state or a series of multiple cognitive states that proceed comparably freely due to a lack of focus and restrictions on the subject matter of any mental state or on the transition between any of the two subsequent mental states [16]. Specifically, mind wandering is considered as a special case of spontaneous thought that is less intentionally restricted than creative thought and objective-directed thinking but more intentionally restricted than dreaming. Another study ( [22]) associated mind wandering with the independence of experiences from impressions and continuing activities, and highlighted the terms “task unrelated” [23] and “stimulus independent” [24] to represent thoughts that occur during mind wandering.

### Workload vs. mind wandering

There have been research efforts focused on understanding the relationship between workload and mind wandering. Zhang *et al.* suggested that there is a negative correlation between mental workload and mind wandering during driving. Specifically, participants with lower cognitive load have additional mental capacity which may be assigned to other tasks irrelevant to the primary task and cause higher mind wandering [25]. Another study claimed that any individual has a fixed mental capacity, and thus, increased perceptual load would lead to diminished available mental capacity for task-irrelevant activities [26]. This is consistent with our hypothesis that higher mind wandering will decrease with increased mental load caused by performing multiple activities at a time.

### Sense of Urgency vs. mind wandering

Latinjak *et al.* proposed a model to examine the relationship between mind wandering and goal-assisted, spontaneous thinking [28]. In particular, the author(s) designed a competition task, settled in a laboratory and performed by athletes, to investigate the link between spontaneous thoughts of the participants and task-unrelated mental activity, and found out that there is a negative correlation between goal-assisted thinking and mind wandering. Considering that, by nature, mind wandering develops when an individual’s attention immediately switches from the current task to inner thoughts [5], it is acceptable to hypothesize that higher mind wandering is negatively correlated with higher sense of urgency (consistent with the proposals in [28]).

### Workload vs. sense of urgency

In contrast to the relationship between workload and mind wandering, and sense of urgency and mind wandering, the relationship between workload and sense of urgency is not entirely clear. Even though there are studies (see sub-section “Workload vs. Sense of Urgency” under section “Related Work”) which examined the correlation between workload and sense of urgency, those research efforts were mainly based on subjective sense of urgency ratings and do not provide moment-by-moment analyses of the impact of sense of urgency occurrences on overall workload.

One important aim of this paper thus was to further investigate the relationship between sense of urgency and overall workload. For one, there will be situations where occasional “sense of urgency moments” do not affect overall task-dependent workload at all. Yet, in other cases frequent events triggering a sense of urgency (such as repeated braking events in a driving task) might lead to increased workload over time. But even if we observe higher workload after frequent sense of urgency moments, the increase in workload might not solely be the result of the task-dependent frequent sense of urgency occurrences as other task factors as well as task-irrelevant activities might have contributed to the overall workload. Hence, it is important to consider settings where no additional factors other than sense of urgency moments are a contributing factor to overall workload.

## Background on Inferring Systemic Cognitive States from Physiological Measures

It is important to note at the outset that past research on using eye gaze, EEG, fNRIS, skin conductance, and respiration for assessing cognitive workload, sense of urgency and mind wandering fundamentally lacks a uniform approach. To the best of our knowledge, no experimental study has examined all three systemic cognitive states, and none has assessed a single systemic cognitive state using all five physiological measures. As a result, the findings are challenging to compare across different experimental paradigms and are often mutually inconsistent. A central aim of this paper is thus to assess the potential for inferring all three systemic cognitive states using all five phyisological measures collected from the same experiment which is possible with the employed dataset [15]. We this were able to provide the first of its kind comprehensive analysis and machine learning effort for isolating the most promising physiological measures for systemtic cognitive state inference in a consistent manner.

### Eye Gaze

#### Eye gaze and workload

Multiple past studies considered pupil diameter as an indicator of cognitive workload [29–33]. One of them recorded eye gaze and EEG from fifteen students in a virtual learning platform for the distinction of cognitive workload levels in two separate groups: students who were able to complete the tasks successfully versus students who were not [29]. The authors found k-Nearest Neighbor as the best predictor over other classification methods. In contrast, we used state-of-the-art classification methods and also utilized neural network-based deep learning models to validate the efficiency of eye gaze in estimating workload. Pfleging *et. al.* introduced a method to assess an individual’s mental workload by using pupil diameter under different lighting situations [33]. Even though the authors proposed a paradigm to estimate cognitive workload using pupillometry, particularly pupil diameter, the small number of participants limits the generalizability of the reported pupil diameter–based workload estimation results.

#### Eye gaze and sense of urgency

There are only a few research papers that explored the efficiency of different physiological signals for determining a sense of urgency. Khalaf *et al.* proposed a method to estimate the participants’ challenge and threat states when they proceed with three mental arithmetic tasks by leveraging different physiological signal types including electrocardiogram (ECG), continuous blood pressure, respiration, impedance cardiogram, and facial electromyography (EMG) [36]. Salkevicius *et al.* studied the assessment of participants’ anxiety levels by leveraging blood volume pressure, galvanic skin response, and skin temperature [37]. To the best of our knowledge, there is no research effort that focuses on understanding the dynamics of human gaze parameters in estimating the sense of urgency levels.

#### Eye gaze and mind wandering

Multiple research efforts have investigated the efficiency of the human gaze in assessing mind wandering [38–40, 90–92, 96, 97]. Bixler *et al.* [38] examined the combination of eye gaze and physiological features to predict mind wandering when the participants were reading instructional texts via the computer interface. He *et al.* [39] examined the efficiency of eye gaze parameters including percentage gaze dwell time spent in the side mirrors and mean of the standard deviation of horizontal gaze position collected from eighteen subjects in estimating mind wandering, their dataset has an inadequate number of participants which might cause their model to overfit. Klesel *et al.* [40] proposed a study to investigate mind wandering in a virtual reality environment by using eye gaze features including fixation coin and fixation duration, but did not provide any experimental finding which validates the efficiency of the human gaze in detecting mind wandering.

### EEG

#### EEG and workload

EEG has been one of the most common physiological markers in predicting human cognitive workload [8, 41–44]. In [43], the authors explored the effects of different frontal lobe frequency bands on assessing working memory workload and observed that decreased delta power was substantially linked to increased workload. Another study defined a workload index, representing the difficulty levels, by leveraging theta and alpha power spectra of EEG found that increased workload was associated with decreased alpha band power and increased theta band power in parietal areas and in prefrontal areas, respectively [44]. In [9], Berka *et al.* investigated the link between task engagement and human workload while completing learning and memory tasks by using EEG signals acquired from eighty healthy participants. Another study first utilized independent component analysis (ICA) to collect the independent features from EEG streams, then estimated the cognitive workload based on these features [45]. Even though we can determine EEG signals directly from the electrical exertion of cortical and sub-cortical neurons with higher temporal resolution, usually in milliseconds, the EEG streams are non-stationary and sensitive to various types of noise including blinking, frequency intrusion, and sensor-based interference associated with motion [46].

#### EEG and sense of urgency

One study investigated the proficiency of EEG and fNIRS in assessing sense of urgency levels by designing a dynamic decision-making setup based on presenting participants with neutral to happy or sad faces. They found out that EEG is able to distinguish between sense of urgency levels which varies person to person [20]. Another research effort explored the effect of long-term (9 seconds) and short-term (5 seconds) time pressure on cognitive responses using mental arithmetic tasks, observing increased arousal levels under higher time pressure based on the analyses of EEG signals [47].

#### EEG and mind wandering

A few studies explored the efficacy of EEG in assessing mind wandering. In [48], Jin *et al.* developed classifiers to examine the state of mind wandering (in particular, either mind-wandering or on-task) by using EEG markers and were able to separate two mind wandering levels with an average accuracy of 60%. Another study analyzed the association between mind wandering and online education by using the spectral power of EEG signals and event-related potential (ERP) waveforms [49]. Dong *et al.* proposed a method for differentiating attention states of participants for both person-dependent and person-independent situations by leveraging ERPs acquired from the EEG signals and concluded that either non-linear or linear ML models can be used to determine mind wandering with the AUC level of 0.715 [50].

### fNIRS

#### fNIRS and workload

Some studies investgiated the dynamics and capabilities of fNIRS signals in assessing different workload levels. Midha *et al.* explored the effectiveness of fNIRS signals, acquired from 20 healthy subjects (8 females and 12 males), in differentiating various human cognitive levels in the case of common office-like tasks including different levels of reading and writing tasks, and observed higher oxygenated haemoglobin (O2Hb) and lower deoxygenated haemoglobin (HHb) in the case of harder reading tasks [53]. In [54], Putze *et al.* examined the performance of fNIRS signals in automatic cognitive workload identification in a virtual reality (VR) setting and suggested that fNIRS signals were capable of detecting mental workload in immensely engaging VR tasks. [52] proposed a method which combines the bi-variate functional brain connectivity (FBC) characteristics of three EEG frequency bands (delta, theta, and alpha) and two fNIRS features (oxyhemoglobin and deoxyhemoglobin) to assess multiple mental workload levels. They concluded that EEG and fNIRS have superior performance in different regions of the brain, *e.g.*, fNIRS performed better in the right frontal region (AF8).

#### fNIRS and sense of urgency

Holtzer *et al.* explored the impact of gender and perceived stress, alone or combined, in pre-frontal cortex (PFC) oxygenated hemoglobin (HBO_2_) variation in the course of walking and observed that perceived stress impacted the task cost, which was specifically observed by HBO_2_ levels of older men [55]. Mirelman *et al.* investigated the fluctuations in brain activity during the primary walking task while performing secondary cognitive tasks by using fNIRS and showed that HBO_2_ can be used to distinguish different stress levels including only walking or walking and counting together [56].

#### fNIRS and mind wandering

There are relatively few efforts focused on assessing mind wandering using fNIRS signals. One study investigated different ML and deep learning classifiers such as deep neural networks (DNNs), convolution neural networks (CNNs), and XGBoost to identify different mind wandering levels without interrupting the incoming task [58]. Another study analyzed the participants’ mind wandering levels by exploring when the participants leave the primary task in the Sustained Attention to Response Task (SART) [59]. They acquired sixteen-channel fNIRS data from frontal cortices and observed increased activity in the area of the medial prefrontal cortex during the mind wandering episodes.

### Skin Conductance

#### Skin conductance and workload

In one study, the authors analyzed the frequency domain features of skin conductance for evaluating mental workload [60]. They modeled a mental task which included memorizing target letters and identifying them from distributed alphabet, and concluded that the frequency domain characteristics of skin conductance were able to detect small changes in cognitive workload. Lew *et al.* proposed a method to determine mental workload from a combination of skin conductance and pupil diameter by using wavelets and genetic programming (GP) and found out that GP had better performance in detecting task performance and difficulty compared to traditional linear discriminant analysis (LDA) [61]. [62] introduced a technique to examine the effectiveness of skin conductance, heart rate, and respiration, acquired from 121 young adults, for predicting mental workload in a simulated driving setting and found that skin conductance was increased with higher task demand.

#### Skin conductance and sense of urgency

Kurniawan *et al.* explored how to measure the change in galvanic skin response (GSR) during acute stress conditions by using multiple classification techniques and found out that in the case of three different workload settings (recovery, light, and heavy), SVM had the highest classification accuracy among four classification methods in differentiating different stress levels [63]. Bakker *et al.* examined GSR data, collected in real-world settings, for assessing stress and argued that the GSR data was ambiguous and further information was needed to make a confident determination of stress levels in addition to GSR signals [64].

#### Skin conductance and mind wandering

We have found comparatively few studies which considered skin conductance as an indicator of different mind wandering levels. Blanchard et. al used the fusion of skin conductance and skin temperature to determine the degree of mind wandering by training various supervised classification methods [65]. Bixler et. al combined eye gaze, skin conductance, and skin temperature along with the contextual information including task difficulty and time on task to assess automatic multimodal detection of mind wandering [38].

### Respiration

#### Respiration and workload

One study examined the association between multiple respiration features (such as inductance plethysmography and pressure of end-tidal carbon dioxide) and different workload levels in simulated flight setting [66]. Specifically, the authors used the NASA-TLX setup and collected respiration signals from seven pilots during one-session simulation test. They found that ventilation has higher values in all tasks compared to baseline. Another study analyzed the performance of multiple physiological signal modalities including heart rate, blood pressure, respiration, and human gaze for assessing mental workload. The authors concluded that the respiratory cycle duration was sensitive to the fluctuations in cognitive workload [67].

#### Respiration and sense of urgency

McDuff *et al.* designed a model to capture cognitive stress via a digital camera by leveraging different physiological measures including breathing rate and heart rate variability, and found that respiration rate increases with increased stress level [68]. Another research study investigated the effects of stress on multiple physiological signal indicators such as skin temperature, facial temperature, and respiration rate, and demonstrated that respiration rate rises with higher stress levels [69].

#### Respiration and mind wandering

In one research work, the authors introduced a method which utilized the temporal synchronization between the respiration signals and the fingertip force pulses, acquired from 12 participants, to measure an objective index to determine mind wandering and suggested that the proposed objective index is a reliable indicator for detecting mind wandering [70]. Another study defined four cognitive state intervals that include mind wandering, awareness of mind wandering, shift of attention, and sustained attention, and explored the performance of respiration rate in monitoring these states and found that respiration rate had no influence in separating mind wandering levels [71].

### Workload vs. Sense of Urgency

There have been a few studies that attempted to characterize the relationship between cognitive workload and the sense of urgency. Wei *et al.* investigated the subjective mental workload of twenty-two residents while entering prescriptions in a computerized physician order entry system by the National Aeronautics and Space Administration-Task Load Index (NASA-TLX) setup and suggested that workload tends to increase with increased sense of urgency level [72]. Keunecke *et al.* related the overall workload to emergency and non-emergency medical transport based on subjective sense of urgency ratings and proposed that workload increases with a higher sense of urgency rating [73]. However, there has been no research effort that investigated the sense of urgency and cognitive workload with the help of eye gaze.

## Methods

We start with a presentation of the employed experimental dataset and also review the definitions of the three systemic cognitive states we set out to explore for which we then define different activation levels based on different experimental conditions. We also review the comprehensive set of physiological signals used to train our computational models.

### Experimental Data and Participants

The utilized dataset [15] consists of multi-modal performance data including pupillometry, electroencephalography (EEG), functional near-infrared spectroscopy (fNIRS), skin conductance, and respiration rate from from 82 participants with the average age of 20 years performing simulated driving tasks. The experimental data were accessed for research purposes beginning on July 1, 2021. Participants were instructed to perform accident-free driving in a realistic driving simulator while completing different secondary tasks. They had to complete two driving sessions, which we refer to as “Session 1” and “Session 2” respectively, with a short break in between. For both sessions, participants were asked to drive on a four-lane highway (two-lane per direction) with a speed limit of 65 mph. During the first three minutes of each session, the only task was driving on a straight road to get participants ready for the rest of the experiment where two other tasks were added:

1. Participants engaged in brief *communication tasks* at different times during each session which included 40 questions in total (20 questions per session). Participants were instructed to respond to the questions, either with “yes/no” and with a longer answer depending on the question type, approximately every 30 to 60 seconds.
2. Participants were forced to brake to avoid accidents about ten times per session during both sessions. In those “braking event”, a front vehicle first came 200 m into the sight of the driver. Next, the driver reached that front vehicle until it was 75 m away and then followed it for 20 seconds at a constant distance of 75 m. Then, the front vehicle immediately decelerated for five seconds with activated brake lights. Finally, the front car accelerated again and moved away from the vehicle (see Figure 2).

**Fig 2.**
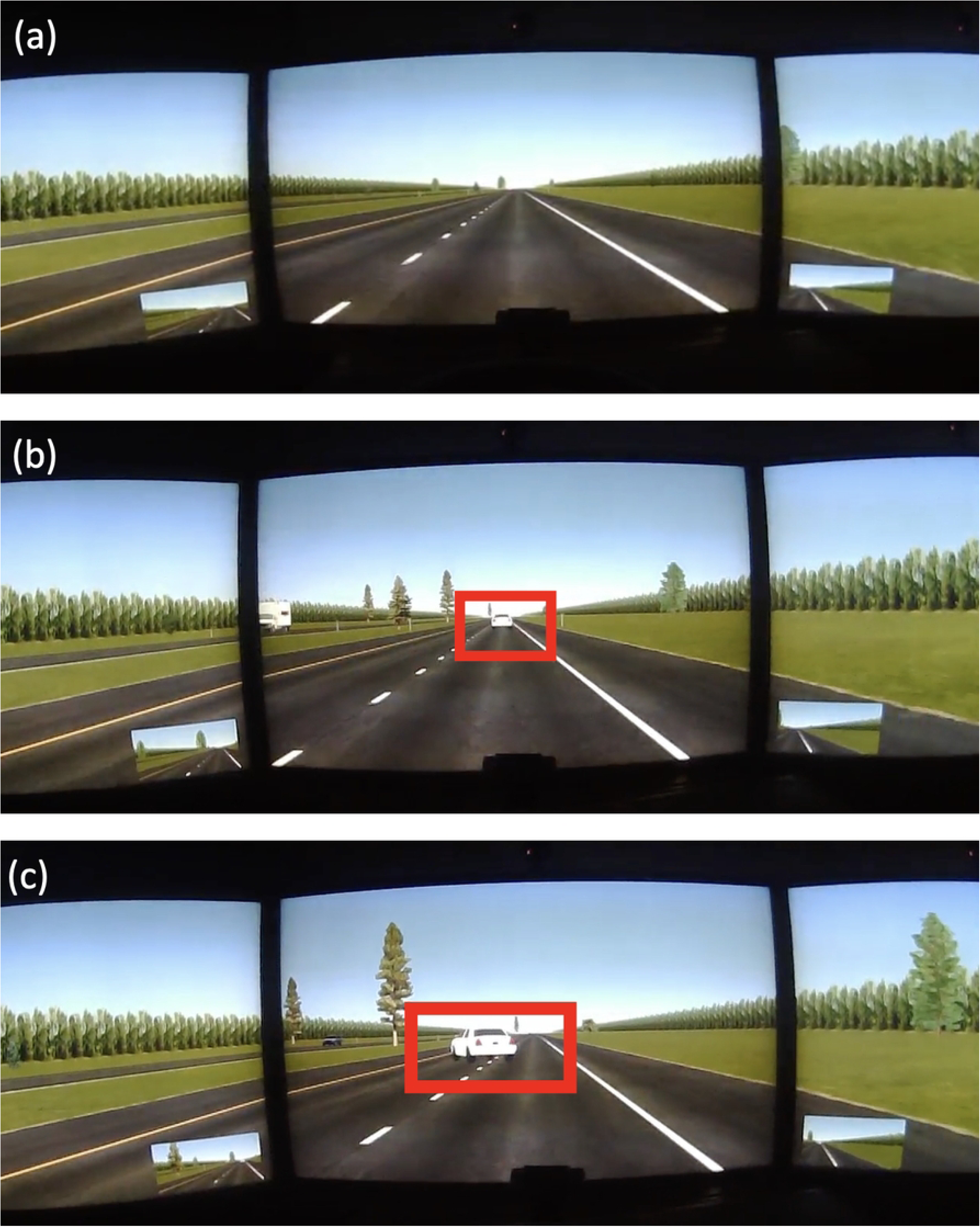
Illustration of a braking event. Example of a braking event: (a) a participant drives in the right lane on a straight road, (b) a car appears in front of the participant’s car, and (c) the front car immediately decelerates, then moves far away.

### Levels of the Systemic Cognitive States

We consider different episodes during the experimental runs for all three systemic cognitive states for which we hypothesized to see different activations of those states (increased or decreased) based on the experimental manipulations that were specifically intended to elicit different levels (see [10] for experimental details). For example, the co-occurrence of a braking event to avoid a crash while performing dialogue events, which requires subjects to answer a question, will lead to competition for attention and thus is likely to increase workload, everything else being equal. In the following sections, we will use the term “level” to refer to those episodes defined by specific experimental manipulations and use the terms “higher” or “lower” to indicate expectations based on the experimental manipulations.

#### Workload Levels

For cognitive workload we define three levels: l_w1_, l_w2_, l_w0_ as shown in Figure 3. Specifically, l_w1_ and l_w2_ start with the 1^st^ and the 5^th^ braking events, respectively, and include four braking and six communication events each. In other words, we defined l_w1_ and l_w2_ by taking different segments during the driving simulation as each of them includes multiple consecutive braking events and dialogue tasks. Here, the onset of braking events corresponds to the appearance of the car in the vehicle’s path. l_w0_ represents the initial period of the driving simulation before the start of the first braking event. We balanced the dataset with a total of 276 samples (92 samples per cognitive workload level) taken from both sessions of 46 subjects.

**Fig 3.**
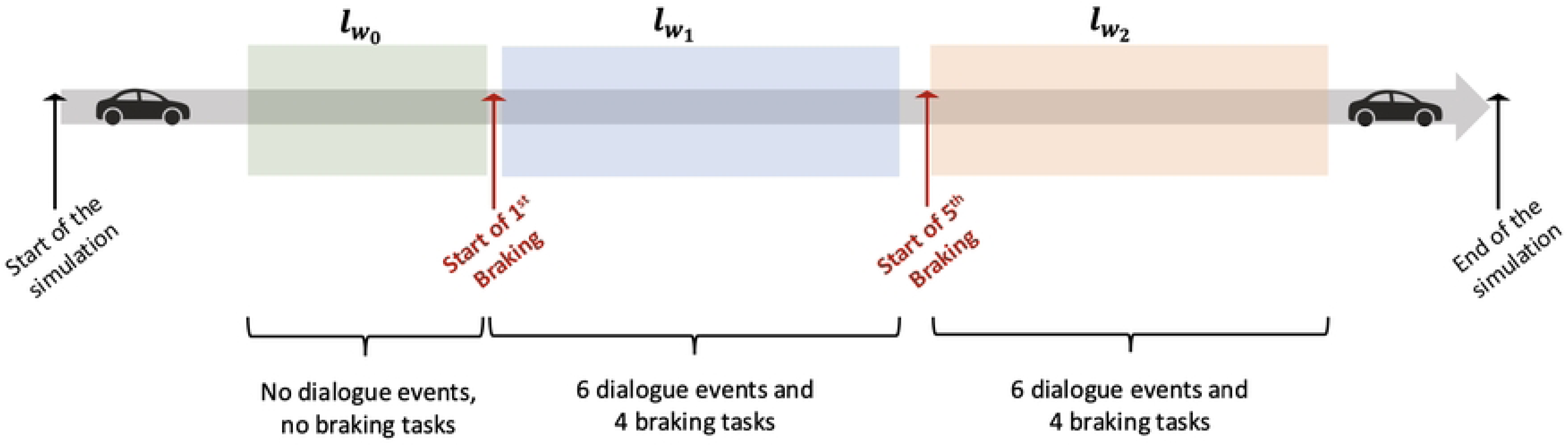
Evaluation of three cognitive workload levels for one session. By selecting different combinations of braking and dialogue events, three different cognitive workload levels were defined.

#### Sense of Urgency Levels

We defined two sense of urgency levels: l_sou1_ and l_sou2_. Specifically, we used the first 2.5 seconds of the 1*^st^* braking event to generate l_sou_ (refer to Figure S4 Fig. Prior experimental work has employed short, event-locked time frames to capture immediate mental effort responses. For example, in a dual-task study of task-evoked responses and effort, participants were given 2.5 seconds to respond to each trial before feedback was provided, explicitly defining the temporal window over which mental effort was elicited and assessed. In line with such experimental paradigms, we used 2.5 seconds windows to capture the rapid, event-locked sense of urgency elicited by sudden events, rather than longer-term workload accumulation [87]. Then, we defined l_sou1_ by taking 2.5 seconds time frames, obtained from 30 seconds before the 1*^st^* braking event to ensure that it does not include any secondary events except from the primary driving task which requires the participants to react rapidly. The rationale is that circumstantial conditions or task demands might require the participants to react immediately and thus generate a higher sense of urgency such as completing a dialog task on time or having to break suddenly during driving [10].

**Fig 4.**
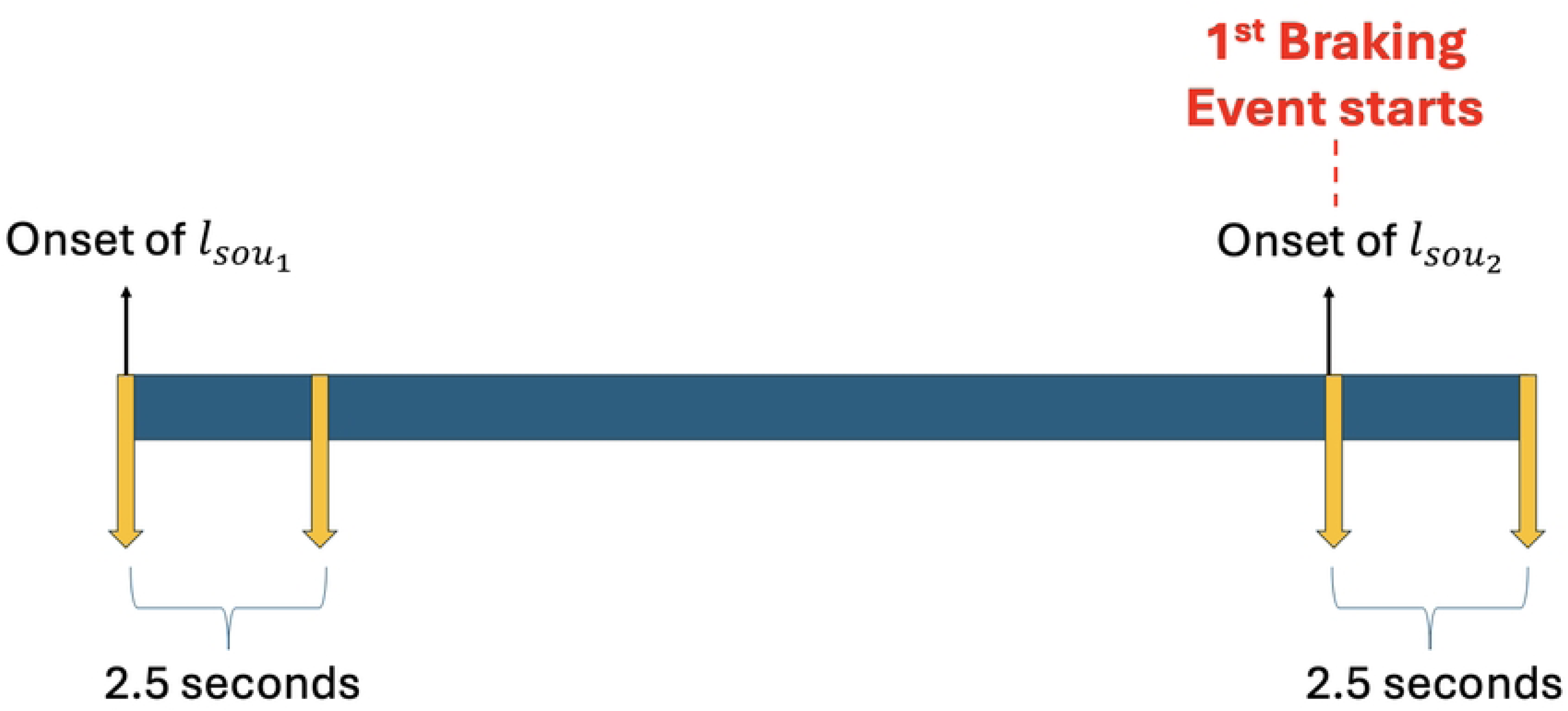
Generation of the two sense of urgency levels. We defined two sense of urgency levels by depicting the onset points of each of them.

#### Mind Wandering Levels

Most of the research works evaluated mind wandering subjectively by using probes in which the participants scored their mind wandering levels themselves during the experiment [99–101]. In contrast, our aim was to design a mind wandering prediction scenario where we could assess mind wandering objectively. For this purpose, we defined two mind wandering levels based on the first 1.5 minutes and the next 1.5 minutes of the simulation, the idea being that during the first 1.5 minutes, the participants were attempting to acclimate to the driving environment which requires initially more attention, resulting in a lower level of mind wandering which we called the “low mind wandering level” l_mw1_ compared to the next 1.5 minutes when participants were already familiar with the driving task. During that time period subjects only had to drive straight on an empty road with no other tasks or distractions, thus requiring the least amount of attention and potentially evoking higher level of mind wandering which we defined as the “high mind wandering level” l_mw2_).

#### Workload vs Sense of Urgency

We also explored how consecutive triggers of the sense of urgency can affect overall workload. In particular, we considered the average PCPS (APCPS) signal during level l_w1_ of a non-DRT session which included multiple subsequent braking tasks and a couple of dialogue interactions. Then, we annotated the onsets of dialogue and braking events and explored the impacts of secondary tasks on the APCPS signal. We conducted a detailed analysis of the signal spikes at the onset points of braking and dialogue events during level l_w1_ of session 1. We hypothesized that these onset points exhibit heightened spikes, which would reflect an increaase in sense of urgency associated with the braking and dialogue tasks during this phase. We obtained a random participant and used their non-DRT session to isolate only braking and dialogue events. We further proposed that the consecutive occurrence of heightened sense of urgency moments might lead to an increase in overall workload.

### Physiological Signal Modalities

We utilized five physiological signal types, including pupillometry, EEG, fNIRS, skin conductance, and respiration, to assess the three different systemic cognitive states. Table 2 shows the characteristics of each signal used in this study including the sampling rate, the pre-processing steps, and the morphological features. We summarized all abbreviations used for these signals in Table 1.

**Table 1.** Notations.

| Term | Description |
| --- | --- |
| PCPS | The percentage change in pupil size |
| PCPS<sub>i</sub> | The PCPS of the i<sup>th</sup> sample |
| APCPS | Average PCPS |
| FixNum | The number of fixations |
| FixDur | The fixation duration |
| BlinkNum | The number of blinks |
| BlinkDur | The blink duration |
| CM | The current measure of pupil diameter |
| BM | The baseline measure of pupil diameter |
| N | The total number of samples in time domain |
| BR | Breathing rate |
| BR<sub>coeff</sub> | The coefficient of variation of BR |
| IBI | The inter-breath interval |
| DC | Duty cycle |
| DC<sub>coeff</sub> | The coefficient of variation of DC |
| IT | The inspiration time |
| ET | The expiration period |
| IE | The ratio between sequential IT and ET |
| ABP | Arterial blood pressure |
| HR | Heart rate |
| IBI | Inter-beat interval |
| SP | Systolic peak |
| DP | Diastolic peak |
| NN interval | Normal-to-normal interval |

**Table 2.** Summary of the employed physiological signal modalities including the sampling rate, the pre-processing steps, and the extracted morphological features.

| Signal Modality | Sampling Rate (Hz) | Pre-processing Steps | Morphological Features |
| --- | --- | --- | --- |
| Pupillometry | 400 | **1-** Amplitude thresholding between 0.8-10 mm<br>**2-** Linear interpolation<br>**3-** 5<sup>th</sup> order Butterworth low-pass filter (LPF) with cutoff frequency of 10 Hz | **1-** Percentage change in pupil size (PCPS)<br>**2-** Average PCPS (APCPS)<br>**3-** Fixation number (per second)<br>**4-** Mean fixation duration<br>**5-** Blink number (per second)<br>**6-** Mean blink duration |
| EEG | 500 | **1-** 6<sup>th</sup> order Butterworth band-pass filter (BPF) between 0.1 Hz - 10 Hz<br>**2-** Independent Component Analysis (ICA)<br>**3-** Kalman Smoother | **1-** Delta<br>**2-** Theta<br>**3-** Alpha<br>**4-** Beta<br>**5-** Gamma |
| fNIRS | 20 | **1-** p-chip interpolation with a sampling rate of 20 Hz<br>**2-** Linear-wise de-trending to find regions in which the variance of the signal is between 25<sup>th</sup> percentile and 75<sup>th</sup> percentile<br>**3-** High-pass filtering with a cutoff frequency of 1.7 Hz<br>**4-** Thresholding to eliminate the channels with the median variance above 1 uM | **1-** The mean of the cleaned channels<br>**2-** The variance of the cleaned channels |
| Skin Conductance | 20 | - | **1-** Mean skin conductance |
| Respiration | 20 | **1-** Upsampling the respiration signal from 20 Hz to 250 Hz<br>**2-** 5<sup>th</sup> order Butterworth low-pass filter (LPF) with cutoff frequency of 1 Hz<br>**3-** Obtaining the local minimas and the local maximas for candidate inhale peaks and exhale troughs<br>**4-** Time thresholding of 0.6 seconds between consecutive inhale peaks and exhale troughs<br>**5-** Amplitude thresholding of 0.2 z-score between consecutive inhale peaks and exhale troughs | **1-** Breathing rate (BR)<br>**2-** Coefficient of variation of BR<br>**3-** Inter-breath interval<br>**4-** Duty cycle (DC)<br>**5-** Coefficient of variation of DC<br>**6-** Inspiration time<br>**7-** Expiration period<br>**8-** I:E |
| Arterial Blood Pressure | 20 | **1-** Upsampling to 250 Hz<br>**2-** 5<sup>th</sup> order Butterworth low-pass filter (BPF) with a cutoff frequency of 5 Hz<br>**3-** Amplitude and beat-to-beat thresholding to identify the systolic pressures (SP) and the diastolic pressures (DP) | **1-** Mean heart rate (HR)<br>**2-** Mean normal-to-normal (NN) interval |

#### Pupillometry

The pupillometry signal was recorded with a sampling rate of 400 Hz. To calculate the pupil diameter, we used the left pupillometry signal considering that the left and the right pupil diameters are synchronous. Next, we applied a three-step pre-processing technique to remove any out-of-band sensory noise and the blink artifact. Particularly, we first utilized amplitude thresholding to eliminate the signal segments below 0.8 mm and above 10 mm by assuming that any values lower than 0.8 mm are potential blink artifacts [74] and the measurable pupil dilation only expands up to at most 10 mm [75]. Secondly, we used linear interpolation to fix the extracted parts [74]. Thirdly, we applied a fifth-order Butterworth low-pass filter with a cutoff frequency of 10 Hz to remove baseline wander [97].

To eliminate subject-based variations in pupil diameter, we calculated the percentage change in pupil size (PCPS) utilizing the following equation [78]:

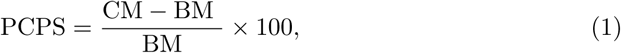

where CM and BM represent the current measure of pupil diameter and the baseline measure of diameter, respectively. Here, BM is determined by calculating the mean of a 10-seconds signal before the stimulus. Additionally, we calculated the average PCPS (APCPS) as follows [78]:

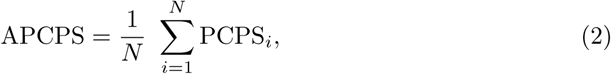

where PCPS*_i_*denotes the percentage change in pupil size at *i^th^* sample and *N* is the total number of samples in the time domain.

#### Electroencephalography

We used three-step pre-processing to denoise the raw EEG signals which were recorded via eight channels with a sampling rate of 500 *Hz*. First, we applied sixth-order Butterworth band-pass filter between 0.1 *Hz* and 30 *Hz* to remove the out-of-band noise [79]. Then, we decomposed the combination of signal epochs to its statistically independent components by leveraging the common technique called independent component analysis (ICA). To manually remove the ICA components which are correlated with blink artifacts, we identified the blink artifacts with the instantaneous spikes in the amplitude. Finally, we applied a Kalman Smoother which is a well-known tool to assess the state of the dynamic linear structures in the presence of noise [80]. We used the Python library called “Tsmoothie” [81] to smooth the EEG signals. We depicted the cross correlation between the EEG signals before and after using Kalman Smoother to show that the smoothing process effectively preserves important features of the original signal (see Fig S5 Fig). Our analyses indicated that the correlation coefficients hover around 0.90 (0.93 and 0.90 for two different signals) which is significantly higher and demonstrated that the essential information contained in the EEG data remains intact, allowing for reliable interpretation of the underlying neural activity.

To statistically analyze the performance of EEG on estimating different human cognitive states, we performed a power spectral density analysis (PSD) and calculated the power distributions of the EEG signals in the frequency domain. Specifically, we obtained the average power of EEG signal from five frequency bands: 1 *Hz* to 4 *Hz* (delta), 4 *Hz* to 8 *Hz* (theta), 8 *Hz* to 12 *Hz* (alpha), 12 *Hz* to 30 *Hz* (beta), 30 *Hz* to 120 *Hz* (gamma), respectively. To obtain the power of these five frequency bands, first we used Welch method to calculate the power spectral density (PSD) which outputs the corresponding frequency and PSD values. Second, we defined the frequency bands as delta (1 Hz to 4 Hz), theta (4 Hz to 8 Hz), alpha (8 Hz to 12 Hz), beta (12 Hz to 30 Hz), and gamma (30 Hz to 120 Hz). Third, we identified the indices for these five frequency bands to determine which frequency values fall within each defined band. Finally, we calculated the absolute power of each frequency band by using Simpson’s rule to approximate the area under the related PSD curve.

**Fig 5.**
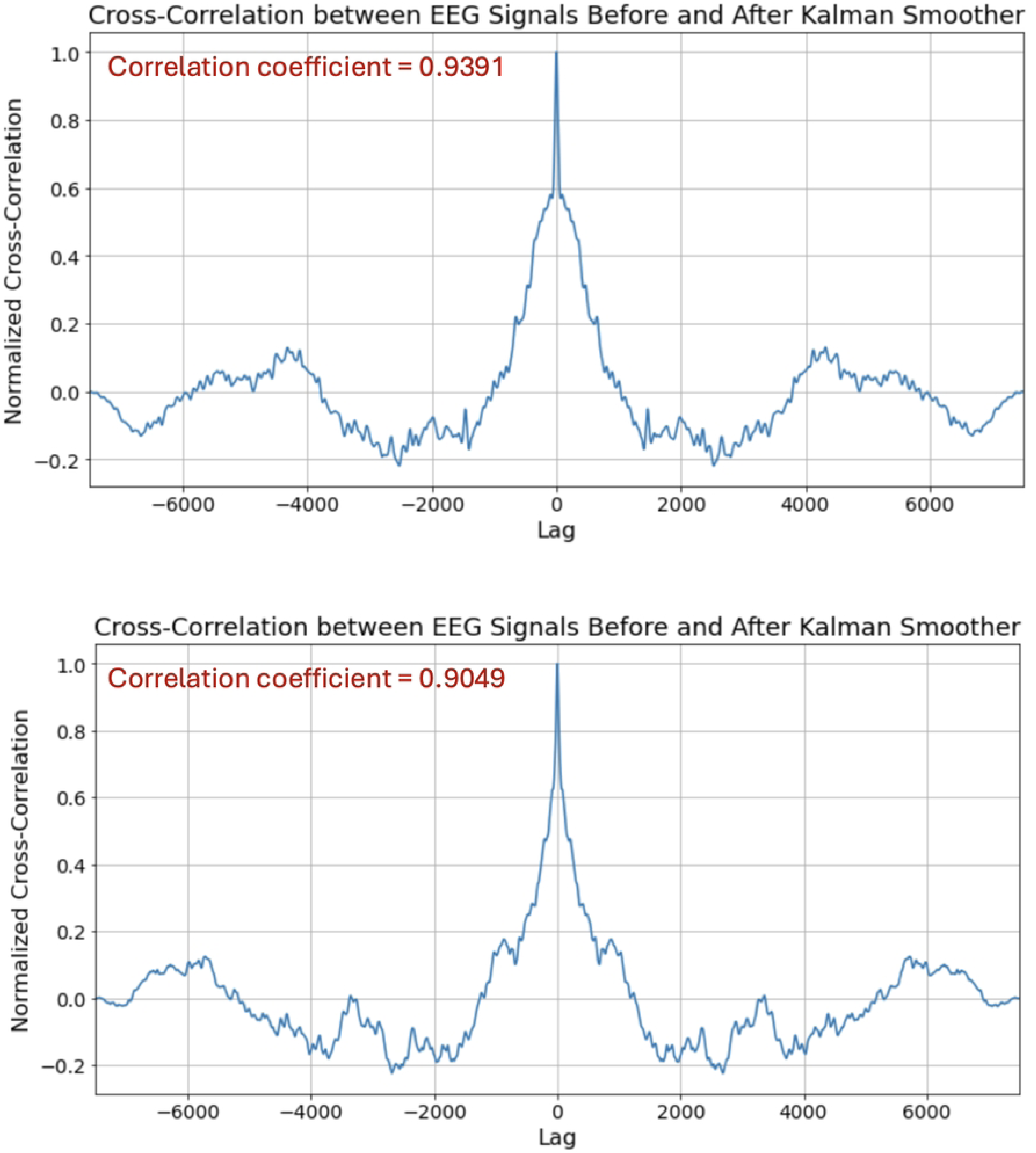
The cross correlation between the EEG signals before and after Kalman Smoother. We depicted the EEG signals before and after applying Kalman Smoother for two different participants.

#### Functional near-infrared spectroscopy

We utilized the cleaning process in [10] to clean the raw fNIRS signal. Next, the mean and variance of fNIRS are extracted which will be used as inputs for learning models in the next sessions.

#### Skin Conductance

Using the skin conductance signal sampled at a rate of 20 Hz, we obtained one well-known and commonly used feature, the mean skin conductance (in microSiemens (muS)), from the raw skin conductance signal.

#### Respiration

We applied five-step pre-processing to denoise the respiration signal which were recorded with a sampling rate of 20 Hz. First, we up-sampled the signal from 20 Hz to 250 Hz. Second, we applied fifth-order Butterworth low-pass filter with a cutoff frequency of 1 *Hz*. Third, we calculated the local minimas and the local maximas to obtain the candidate inhale peaks and exhale troughs. Fourth, we applied time thresholding of 0.6 seconds between the sequential inhale peaks and exhale troughs. Finally, we used amplitude thresholding of 0.2 z-score between the sequential inhale peaks and exhale troughs to acquire the final set of respiration beats. In this study, we collected eight morphological features from the respiration signal [57, 82] which we summarized as follows:

1. **The breathing rate (BR)** (in 1/sec) which represents the time between the inhale onsets.
2. **The coefficient of variation of the breathing rate (BR_coeff_)** (in 1/sec) which is proportion of the standard deviation of the difference between inhale onsets over the average difference between inhale onsets.
3. **The inter-breath interval (IBI)** (in sec) which represents the average time between the inhale onsets.
4. **The duty cycle (DC)** which represents the proportion of breath that is inhaled calculated as the proportion of the average inhale duration over the average inter-breath interval.
5. **The coefficient of variation of duty cycle (DC_coeff_)** which is the proportion of the standard deviation of inhale duration over the average inhale duration.
6. **The I:E (IE)** which represents the ratio between the sequential inspiration time and the expiration period.
7. **The inspiration time (IT)** (in sec) which is the time duration between a trough and a peak within the tidal volume waveform (TVW) signal.
8. **The expiration period (ET)** (in sec) which is the time duration between a peak and a trough within the TVW signal.

#### Arterial Blood Pressure

We also included arterial blood pressure to evaluate the performance of this signal modality for predicting different human cognitive states in comparison to another technique proposed by *Borisov et. al.* [98] (the results are presented in the section named “Comparison the Performance of Pupillometry, Arterial Blood Pressure, and Skin Conductance in Classifying Different Cognitive States”). We pre-processed the ABP signal by following three steps. First, we upsampled the ABP signal from 20 Hz to 250 Hz. Second, we applied a 5*^th^* order Butterworth low-pass filter with a cutoff frequency of 5 *Hz* to eliminate the out-of-band noise. Third, we identified the systolic pressures (SP) and diastolic pressures (DP) by using both an amplitude threshold and a beat-to-beat duration threshold based on the behavior of cardiac dynamics. To accomplish these, we determined the SP and DP candidates by taking the local maxima and local minima spikes from the ABP signal, respectively. We evaluated the SP and DP candidates as the potential heartbeats given that SP points taken from ABP signals are reliable markers to calculate heart rate and interbeat interval (IBI) [93–95]. Here, IBI represents the time duration between two sequential heartbeats. After that, we identified the amplitude threshold, which shows the threshold for the difference between the consecutive DP and SP points, as 10 *mmHg*. Namely, we determined the subsequent DP and SP points whose amplitudes differ more than 10 *mmHg* as candidate SP and DP points. Next, we selected the minimum and the maximum IBI values as 30 and 120 beats per minute (bpm), respectively. The intuition behind this choices was that the common resting HR values of an adult lie within 60 bpm and 90 bpm [17]. We used wider range of HR to prevent the false detection of HR values by considering subject-based variations.

We considered the SP points as the heartbeats and used them to calculate the IBI. We obtained the HR from the estimated IBI by using the following equation:

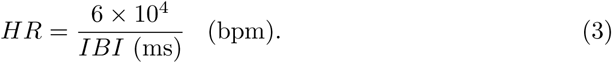

Based on the estimated HR and IBI values, we acquired two biomarkers 1- Mean HR and 2- Mean normal-to-normal (NN) interval.

## Statistical Analyses of Systemic Cognitive State Levels

We performed comprehensive statistical analyses using the five physiological signal modalities to determine the best modalities for detecting different levels of cognitive workload, sense of urgency, and mind wandering. Specifically, we used ANOVA statistical tests with Tukey’s “honestly significant difference” (HSD) multiple pairwise comparisons [84] to investigate the variations of different physiological indicators for three workload levels (l_w0_, l_w1_, and l_w2_), two sense of urgency levels (l_sou1_ and l_sou2_), and two mind wandering levels (l_mw1_ and l_mw2_).

### Workload

Figure S6 Fig shows the boxplot of the mean APCPS taken from both of the sessions, assessing different human cognitive workload levels. The results demonstrate that APCPS is a reliable indicator to separate multiple workload levels. First, we ran full ANOVA models on three workload levels and obtained the *f* -value, the *p*-value, and the partial *η*^2^ by using the APCPS obtained only from session 1, only from session 2, and from both of the sessions which we reported the results in Table 3 (above). The results showed that the three worklaod levels can be distinguishable by utilizing the APCPS biomarker. Then, we proceeded with the post-hoc pairwise comparisons and performed Tukey’s HSD multiple pairwise comparison test to examine the variation in the mean APCPS among the three cognitive workload levels. Table 3 (below) shows the results of the p-values for all pairs of workload levels obtained from Tukey’s HSD multiple pairwise comparison test at a significance level of .95 by using the APCPS. The results demonstrate that there is a statistically significant difference between all pairs of workload levels (p-value *<*.05). Additionally, Figure S7 Fig depicts the mean APCPS values related to separate sessions for different workload levels. The results demonstrate that we can differentiate session 1 and session 2 during l_w1_ by using the APCPS. On the other hand, the difference between the APCPS values of session 1 and session 2 during l_w2_ and l_w0_ are negligible. This suggests a training effect of session 1 during l_w1_ which we required the participants to acclimate with different types of secondary events including braking tasks and dialogue communications.

**Fig 6.**
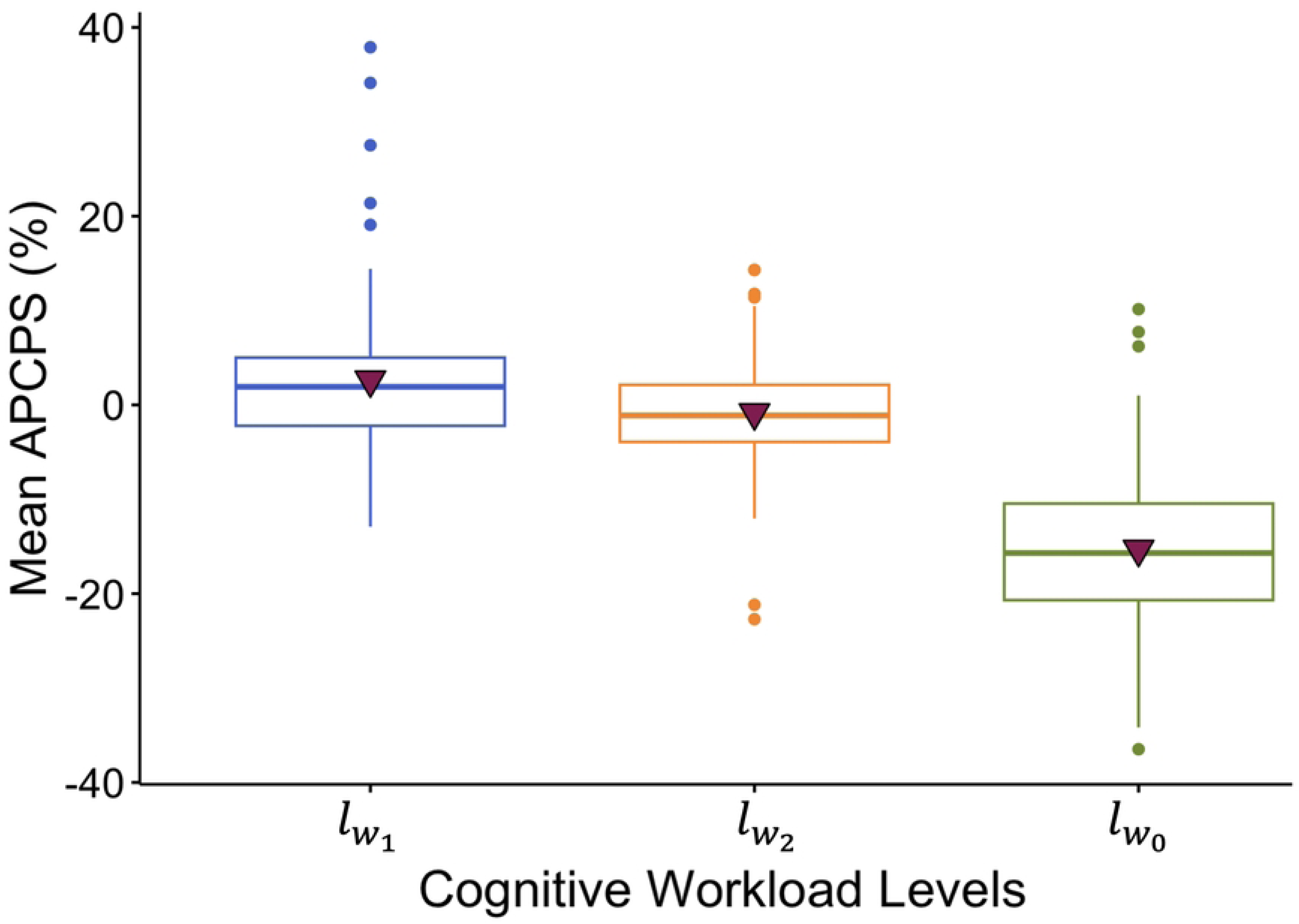
The change in the mean APCPS related to different cognitive workload levels. Boxplot of the mean APCPS values corresponding to different cognitive workload levels including both sessions from all of the subjects.

**Fig 7.**
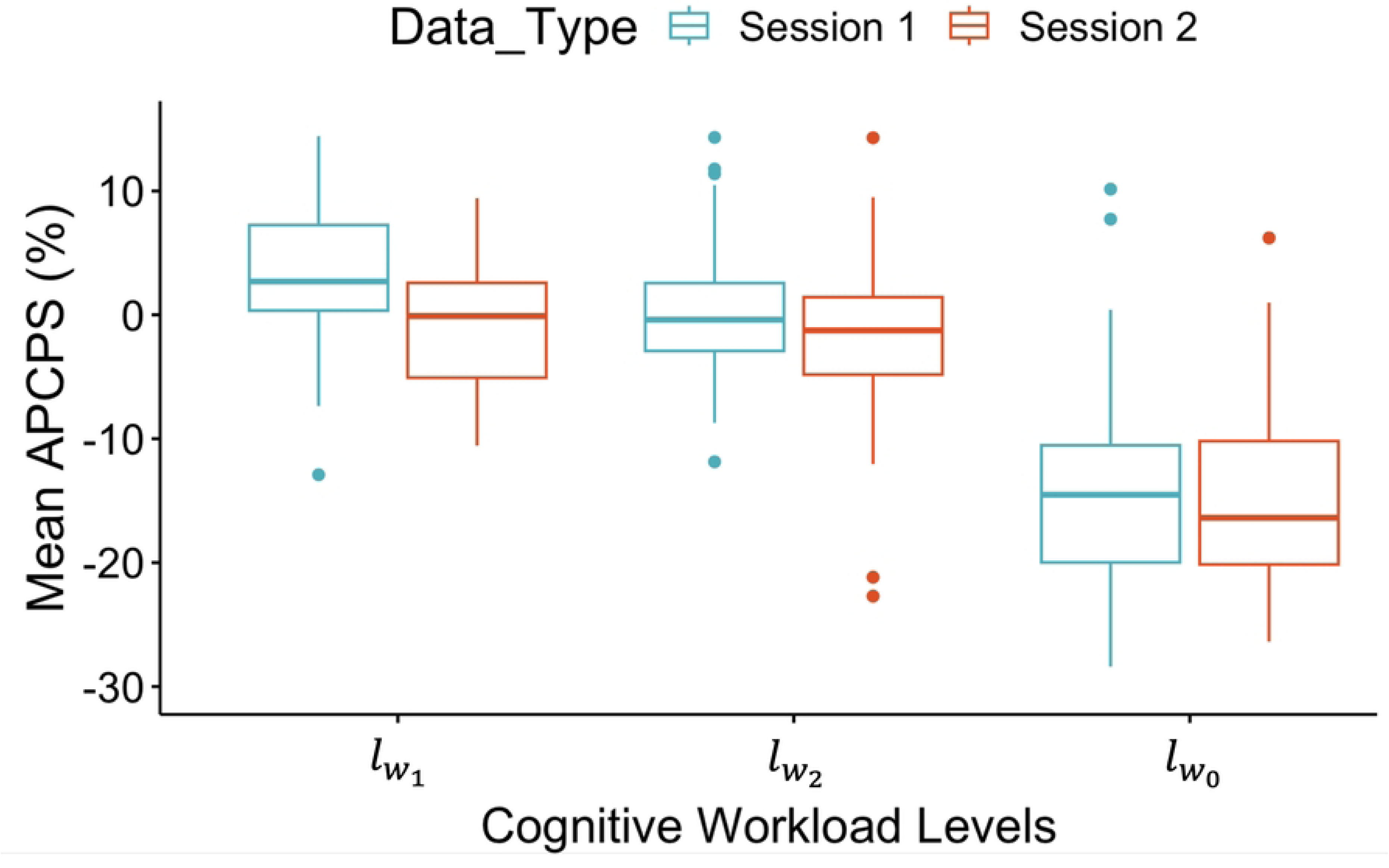
The change in the mean APCPS related to different cognitive workload levels for two sessions. Boxplot of the mean APCPS values corresponding to different sessions of three cognitive workload levels.

**Table 3.** Results of the full ANOVA model and Tukey’s HSD multiple pairwise test for different pairs of cognitive workload levels by using the mean APCPS.

| Full ANOVA model results | f-value | p-value | Partial Eta Squared |
| --- | --- | --- | --- |
| Session 1 | 79.74 | $1.37 \times 10^{-23}$ | 0.54 |
| Session 2 | 63.14 | $4.40 \times 10^{-20}$ | 0.48 |
| In total | 137.91 | $4.02 \times 10^{-42}$ | 0.50 |

| Tukey’s HSD Test | p-value (Session 1) | p-value (Session 2) | p-value (in total) |
| --- | --- | --- | --- |
| $l_{w_1} - l_{w_0}$ | $2.23 \times 10^{-13}$ | 0.00 | $1.76 \times 10^{-14}$ |
| $l_{w_2} - l_{w_0}$ | $2.23 \times 10^{-13}$ | $2.52 \times 10^{-14}$ | $2.75 \times 10^{-14}$ |
| $l_{w_1} - l_{w_2}$ | $7.47 \times 10^{-3}$ | $8.74 \times 10^{-3}$ | $3.81 \times 10^{-1}$ |

To investigate the efficiency of 21 physiological biomarkers in addition to the APCPS, obtained from multiple signal modalities including EEG, fNIRS, skin conductance, and respiration, we ran again full ANOVA models on three workload levels to obtain *f* -value, *p*-value, and partial *η*^2^ which we reported in Table 4. The results show that other than the APCPS, all five EEG bands and two respiration features (*BR_coeff_* and IBI) have significant *p*-values. Then, we proceeded to perform post-hoc multiple pairwise analysis by using Tukey’s HSD multiple pairwise test which we presented the results in Table 5. The results demonstrated that the APCPS is the best feature in classifying all pairs of cognitive workload levels among other physiological features. We observed that, despite the significant *p*-values obtained from full ANOVA model, only one of the EEG features (beta) was able to separate all pairs of workload levels. Despite the *p*-values obtained from EEG (beta) were lower than .05, they were relatively higher than the *p*-values obtained from the APCPS. Even though two respiration features were found to have significant p-values from full ANOVA model test, none of them were able to distinguish all pairs of three workload levels (by using Tukey’s HSD multiple pairwise test). We also reported the mean and the standard deviation values in Table 6.

**Table 4.**
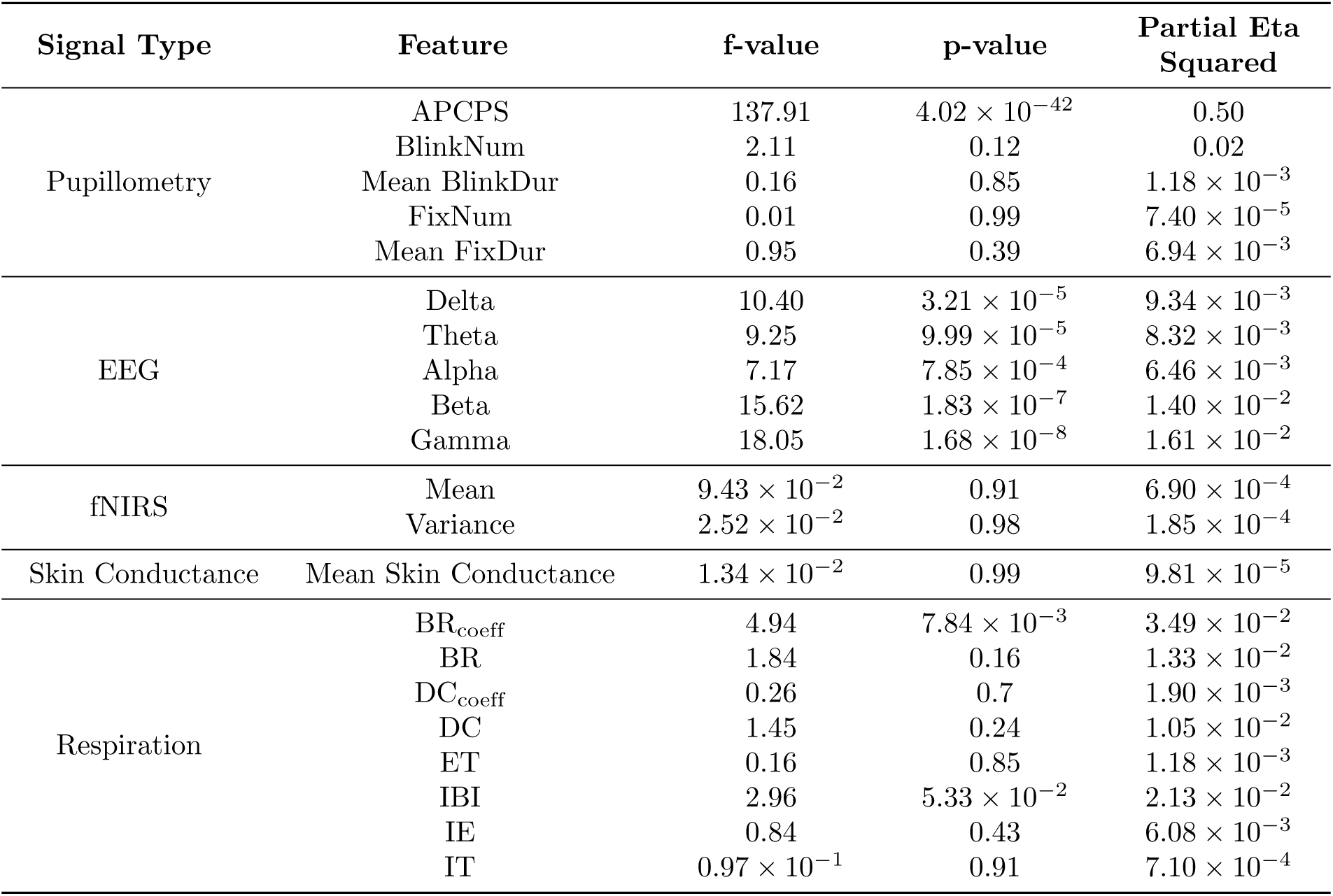
Full ANOVA model results for the morphological features obtained from different physiological signal types including pupillometry, EEG, fNIRS, skin conductance, and respiration for differentiating the cognitive workload levels.

**Table 5.**
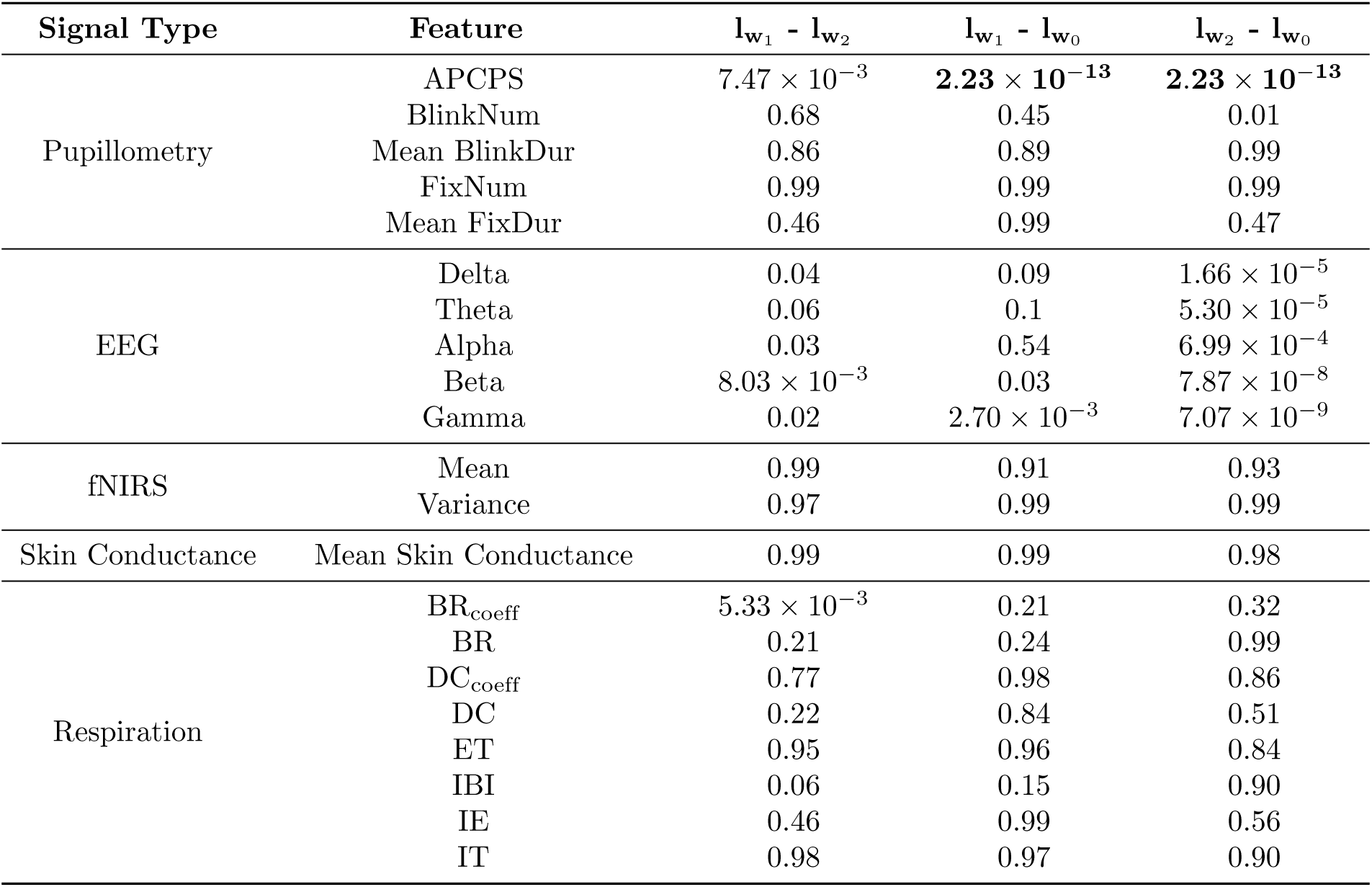
p-values of Tukey HSD multiple pairwise test for the morphological features obtained from different physiological signal types including pupillometry, EEG, fNIRS, skin conductance, and respiration for different pairs of cognitive workload levels.

| Signal Type | Feature | $l_{w_1} - l_{w_2}$ | $l_{w_1} - l_{w_0}$ | $l_{w_2} - l_{w_0}$ |
| --- | --- | --- | --- | --- |
| Pupillometry | APCPS | $7.47 \times 10^{-3}$ | **$2.23 \times 10^{-13}$** | **$2.23 \times 10^{-13}$** |
|  | BlinkNum | 0.68 | 0.45 | 0.01 |
|  | Mean BlinkDur | 0.86 | 0.89 | 0.99 |
|  | FixNum | 0.99 | 0.99 | 0.99 |
|  | Mean FixDur | 0.46 | 0.99 | 0.47 |
| EEG | Delta | 0.04 | 0.09 | $1.66 \times 10^{-5}$ |
| | Theta | 0.06 | 0.1 | $5.30 \times 10^{-5}$ |
| | Alpha | 0.03 | 0.54 | $6.99 \times 10^{-4}$ |
| | Beta | $8.03 \times 10^{-3}$ | 0.03 | $7.87 \times 10^{-8}$ |
| | Gamma | 0.02 | $2.70 \times 10^{-3}$ | $7.07 \times 10^{-9}$ |
| fNIRS | Mean | 0.99 | 0.91 | 0.93 |
|  | Variance | 0.97 | 0.99 | 0.99 |
| Skin Conductance | Mean Skin Conductance | 0.99 | 0.99 | 0.98 |
| Respiration | BR<sub>coeff</sub> | $5.33 \times 10^{-3}$ | 0.21 | 0.32 |
|  | BR | 0.21 | 0.24 | 0.99 |
|  | DC<sub>coeff</sub> | 0.77 | 0.98 | 0.86 |
|  | DC | 0.22 | 0.84 | 0.51 |
|  | ET | 0.95 | 0.96 | 0.84 |
|  | IBI | 0.06 | 0.15 | 0.90 |
|  | IE | 0.46 | 0.99 | 0.56 |
|  | IT | 0.98 | 0.97 | 0.90 |

**Table 6.** The mean and the standard deviation of the morphological features obtained from different physiological signal types including pupillometry, EEG, fNIRS, skin conductance, and respiration for different pairs of cognitive workload levels.

| Signal Type | Feature | (Mean ± SD) of $l_{w_0}$ | (Mean ± SD) of $l_{w_1}$ | (Mean ± SD) of $l_{w_2}$ |
| --- | --- | --- | --- | --- |
| Pupillometry | APCPS | -15.3 ± 8.27 | 2.65 ± 8.53 | -0.83 ± 6.32 |
| | BlinkNum | 0.43 ± 0.22 | 0.47 ± 0.24 | 0.50 ± 0.24 |
| | Mean BlinkDur | 0.39 ± 0.30 | 0.37 ± 0.32 | 0.40 ± 0.39 |
| | FixNum | 2.88 ± 0.76 | 2.88 ± 0.76 | 2.89 ± 0.75 |
| | Mean FixDur | 0.19 ± 0.08 | 0.19 ± 0.08 | 0.18 ± 0.07 |
| EEG | Delta | 1.70 ± 1.22 | 1.88 ± 1.40 | 2.09 ± 2.09 |
| | Theta | 17.7 ± 22.9 | 21.3 ± 26.3 | 25.1 ± 45.6 |
| | Alpha | 0.10 ± 0.09 | 0.11 ± 0.09 | 0.12 ± 0.13 |
| | Beta | $(5.23 \pm 4.26) \times 10^{-3}$ | $(5.95 \pm 4.72) \times 10^{-3}$ | $(6.77 \pm 6.63) \times 10^{-3}$ |
| | Gamma | $(2.29 \pm 0.98) \times 10^{-6}$ | $(3.35 \pm 0.14) \times 10^{-6}$ | $(4.22 \pm 0.16) \times 10^{-6}$ |
| fNIRS | Mean | 0.40 ± 0.06 | 0.39 ± 0.06 | 0.39 ± 0.06 |
| | Variance | 0.08 ± 0.03 | 0.08 ± 0.03 | 0.08 ± 0.03 |
| Skin Conductance | Mean Skin Conductance | 5.52 ± 2.77 | 5.51 ± 2.81 | 5.46 ± 2.75 |
| Respiration | BR<sub>coeff</sub> | 24.1 ± 14.6 | 27.0 ± 12.2 | 30.4 ± 14.1 |
| | BR | 18.8 ± 2.93 | 18.8 ± 2.63 | 18.2 ± 2.45 |
| | DC<sub>coeff</sub> | 26.9 ± 16.9 | 28.0 ± 14.5 | 28.4 ± 12.6 |
| | DC | 0.47 ± 0.14 | 0.45 ± 0.14 | 0.43 ± 0.15 |
| | ET | 3.53 ± 1.77 | 3.37 ± 1.91 | 3.45 ± 1.87 |
| | IBI | 3.43 ± 0.64 | 3.47 ± 0.63 | 3.65 ± 0.70 |
| | IE | 0.53 ± 0.14 | 0.55 ± 0.14 | 0.55 ± 0.13 |
| | IT | 1.59 ± 0.55 | 1.55 ± 0.64 | 1.57 ± 0.63 |

### Sense of Urgency

Figure S8 Fig shows the boxplot of the mean APCPS values corresponding to lower and higher sense of urgency levels. The results demonstrate that APCPS is capable of differentiating two sense of urgency levels. We observed that the APCPS is notably lower for less sense of urgency (l_sou1_) compared to higher sense of urgency levels (l_sou2_).

**Fig 8.**
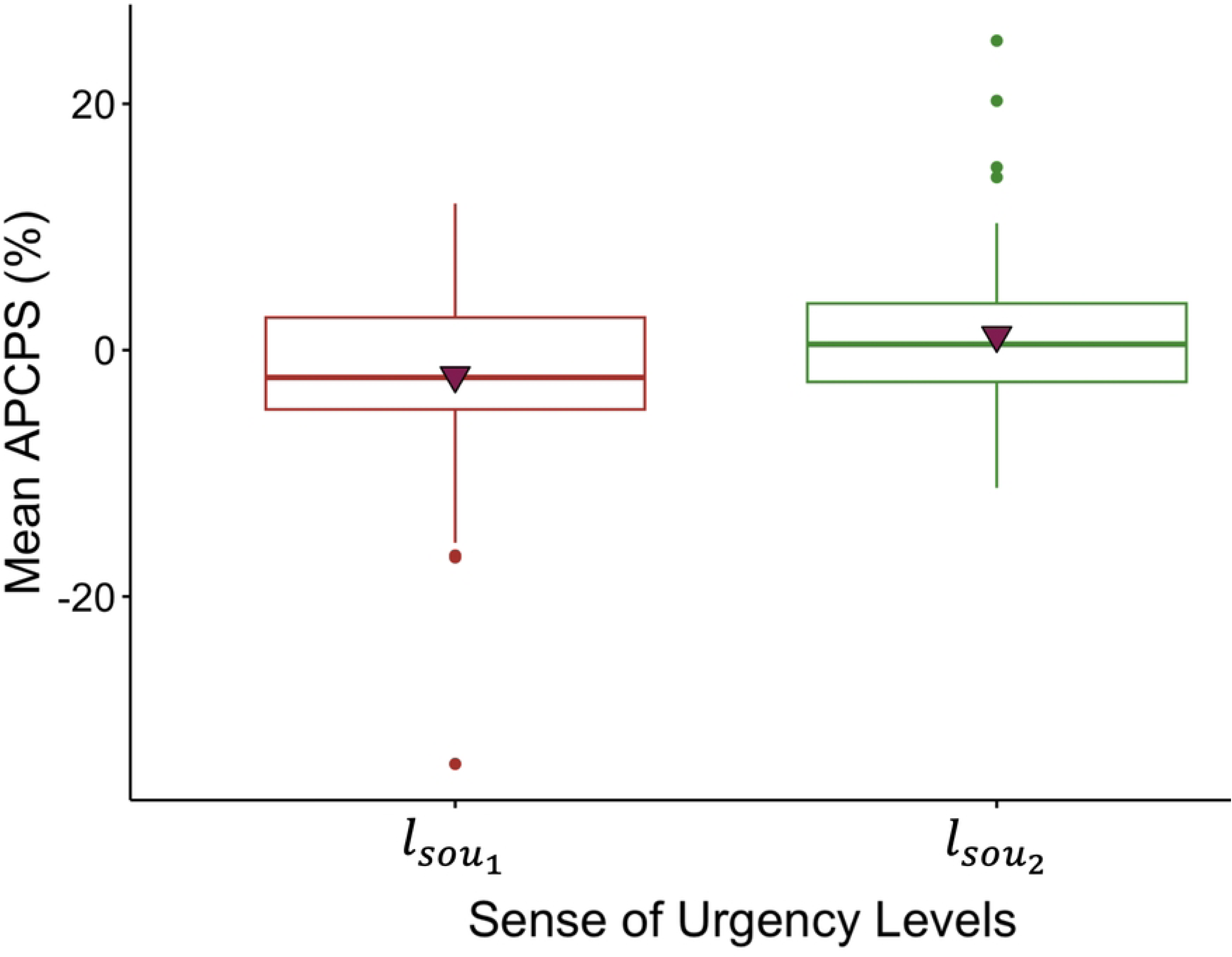
The change in the mean APCPS related to different sense of urgency levels. Boxplot of the mean APCPS corresponding to two sense of urgency levels including both sessions from all of the subjects. l_sou1_: Lower sense of urgency, l_sou2_: Higher sense of urgency.

Furthermore, we explored the effectiveness of other physiological indicators in differentiating two sense of urgency levels. Table 7 indicates (a) *p*-values of Tukey HSD multiple pairwise test and (b) full ANOVA models for 11 physiological markers obtained from four signal types. In addition, we added the mean and the standard deviation of the values to be more informative by indicating how the values of different biomarkers fluctuate in response to changing the sense of urgency levels. Here, we excluded eight respiration (BR_coeff_, BR, DC_coeff_, DC, ET, IBI, IE, and IT) and two eye gaze features (BlinkNum and mean BlinkDur) given that we explored 2.5-seconds signal segment which may not include enough number of respiration or blinking to calculate the related physiological markers. The results indicate that APCPS is the best feature in separating two sense of urgency levels compared to other physiological signal indicators. On the other hand, there were three EEG features (including alpha, beta, and gamma) which were also able to differentiate l_sou1_ and l_sou2_. In particular, the performance of alpha (EEG) was closely equivalent to the performance of the APCPS (pupillometry) which guided us to further investigate the performance of the fusion of pupillometry and EEG in classifying two sense of urgency levels.

**Table 7.**
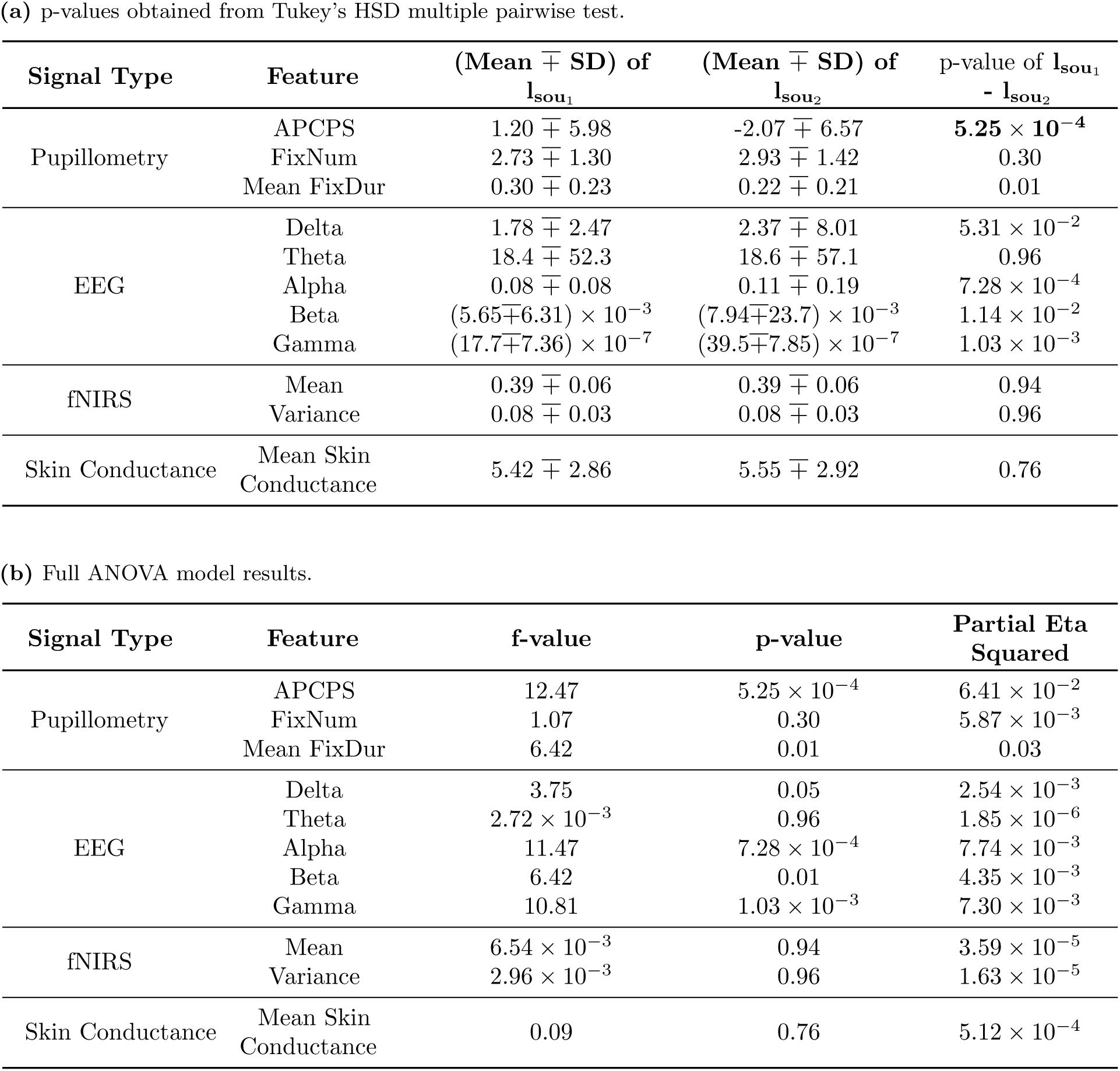
(a) p-values of Tukey HSD multiple pairwise test and (b) full ANOVA model results for the morphological features obtained from different physiological signal types including pupillometry, EEG, fNIRS, and skin conductance for two sense of urgency levels.

| Signal Type | Feature | (Mean ± SD) of $l_{sou_1}$ | (Mean ± SD) of $l_{sou_2}$ | p-value of $l_{sou_1} - l_{sou_2}$ |
| --- | --- | --- | --- | --- |
| Pupillometry | APCPS | $1.20 \pm 5.98$ | $-2.07 \pm 6.57$ | $5.25 \times 10^{-4}$ |
| | FixNum | $2.73 \pm 1.30$ | $2.93 \pm 1.42$ | 0.30 |
| | Mean FixDur | $0.30 \pm 0.23$ | $0.22 \pm 0.21$ | 0.01 |
| EEG | Delta | $1.78 \pm 2.47$ | $2.37 \pm 8.01$ | $5.31 \times 10^{-2}$ |
| | Theta | $18.4 \pm 52.3$ | $18.6 \pm 57.1$ | 0.96 |
| | Alpha | $0.08 \pm 0.08$ | $0.11 \pm 0.19$ | $7.28 \times 10^{-4}$ |
| | Beta | $(5.65 \pm 6.31) \times 10^{-3}$ | $(7.94 \pm 23.7) \times 10^{-3}$ | $1.14 \times 10^{-2}$ |
| | Gamma | $(17.7 \pm 7.36) \times 10^{-7}$ | $(39.5 \pm 7.85) \times 10^{-7}$ | $1.03 \times 10^{-3}$ |
| fNIRS | Mean | $0.39 \pm 0.06$ | $0.39 \pm 0.06$ | 0.94 |
| | Variance | $0.08 \pm 0.03$ | $0.08 \pm 0.03$ | 0.96 |
| Skin Conductance | Mean Skin Conductance | $5.42 \pm 2.86$ | $5.55 \pm 2.92$ | 0.76 |

(b) Full ANOVA model results.
| Signal Type | Feature | f-value | p-value | Partial Eta Squared |
| --- | --- | --- | --- | --- |
| Pupillometry | APCPS | 12.47 | $5.25 \times 10^{-4}$ | $6.41 \times 10^{-2}$ |
| | FixNum | 1.07 | 0.30 | $5.87 \times 10^{-3}$ |
|  | Mean FixDur | 6.42 | 0.01 | 0.03 |
| EEG | Delta | 3.75 | 0.05 | $2.54 \times 10^{-3}$ |
| | Theta | $2.72 \times 10^{-3}$ | 0.96 | $1.85 \times 10^{-6}$ |
| | Alpha | 11.47 | $7.28 \times 10^{-4}$ | $7.74 \times 10^{-3}$ |
| | Beta | 6.42 | 0.01 | $4.35 \times 10^{-3}$ |
| | Gamma | 10.81 | $1.03 \times 10^{-3}$ | $7.30 \times 10^{-3}$ |
| fNIRS | Mean | $6.54 \times 10^{-3}$ | 0.94 | $3.59 \times 10^{-5}$ |
| | Variance | $2.96 \times 10^{-3}$ | 0.96 | $1.63 \times 10^{-5}$ |
| Skin Conductance | Mean Skin Conductance | 0.09 | 0.76 | $5.12 \times 10^{-4}$ |

### Mind Wandering

Figure S9 Fig shows the boxplot of the mean APCPS values related to lower and higher mind wandering levels taken from both of the sessions. The results indicated that mean APCPS is considerably higher in l_mw1_ (less mind wandering) than in l_mw2_ (more mind wandering where the participants were used to driving and were responsible for proceeding only with driving without additional secondary tasks). We also ran Tukey’s HSD pairwise test and found out that the *p*-value is 3.60 × 10^-14^ in differentiating l_mw1_ and l_mw2_ at a significance level of .95. The overall results demonstrated that APCPS is a reliable indicator in predicting different mind wandering levels.

**Fig 9.**
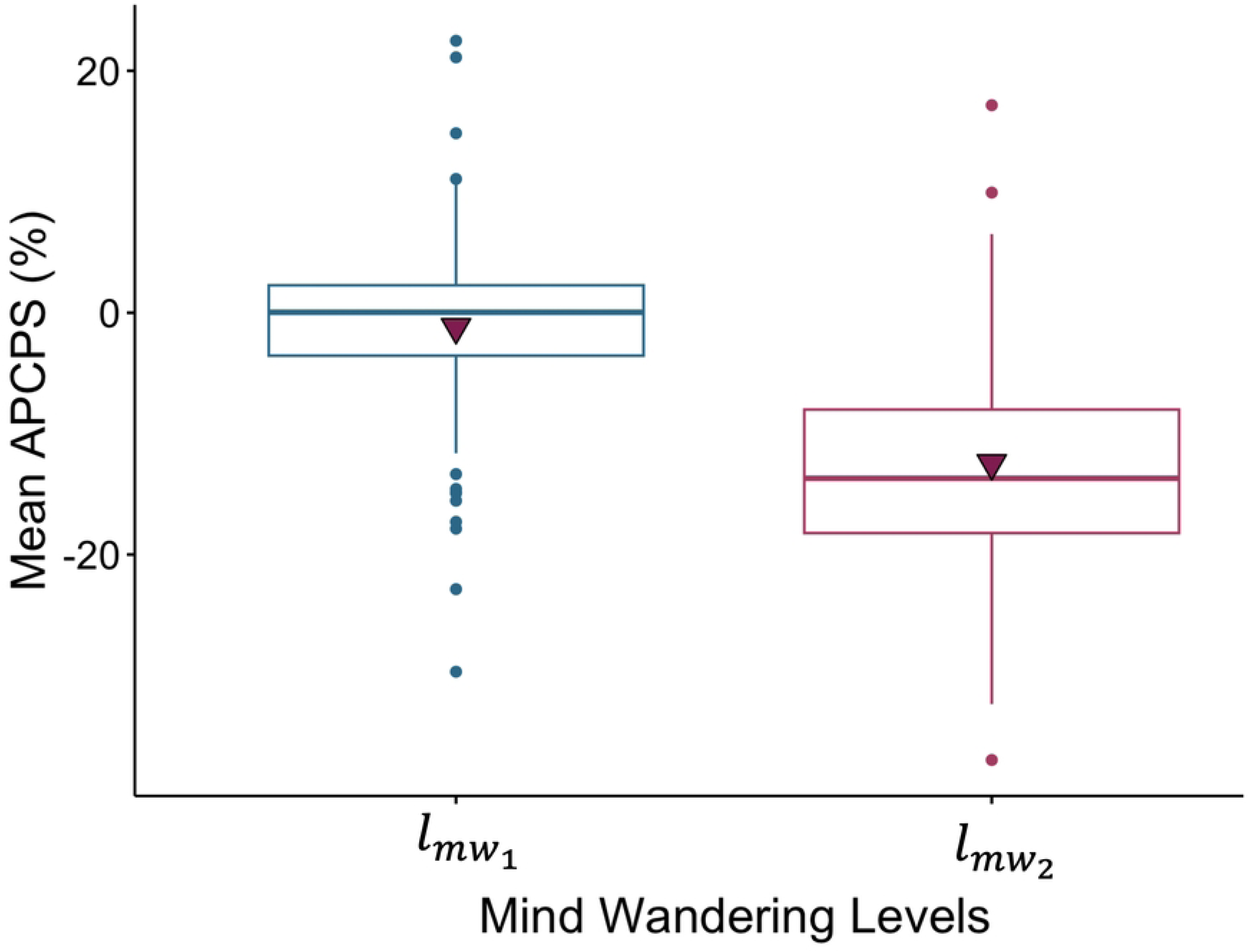
The change in the mean APCPS related to different mind wandering levels. Boxplot of the mean APCPS corresponding to two mind wandering levels including both sessions from all of the subjects. l_mw1_: Lower mind wandering, l_mw2_: Higher mind wandering.

We further analyzed the performance of other physiological metrics in classifying two mind wandering levels in addition to the APCPS. Table 8 (a) shows the *p*-values obtained from Tukey’s HSD pairwise test as well as the mean and the standard deviation values, and Table 8 (b) indicates the results for full ANOVA models including *f*-value, *p*-value, and partial *η*^2^ for 21 physiological markers acquired from five signal types. The *p*-values obtained from Tukey’s HSD multiple pairwise test represent the differentiation performance of two mind wandering levels by using different physiological biomarkers. We tested 21 different physiological features individually to analyze their performance in separating two mind wandering levels. The results show that APCPS is the most effective indicator in differentiating l_mw1_ and l_mw2_ compared to other physiological markers.Even though there were some other indicators including theta and alpha powers obtained from EEG which have p-values lower than .05, they were significantly higher that the *p*-value obtained from the APCPS which makes the APCPS the most reliable physiological feature in assessing different mind wandering levels.

**Table 8.** p-values of Tukey HSD multiple pairwise test as well as the mean and the standard deviation for the morphological features obtained from different physiological signal types including pupillometry, EEG, fNIRS, skin conductance, and respiration for two mind wandering levels.

| Signal Type | Feature | (Mean ± SD) of $l_{mw_1}$ | (Mean ± SD) of $l_{mw_2}$ | p-value of $l_{mw_1} - l_{mw_2}$ |
| --- | --- | --- | --- | --- |
| Pupillometry | APCPS | $-1.19 \pm 0.02$ | $-12.4 \pm 8.85$ | $3.60 \times 10^{-14}$ |
| | BlinkNum | $0.43 \pm 2.22$ | $0.41 \pm 2.22$ | 0.59 |
| | Mean BlinkDur | $0.40 \pm 0.37$ | $0.43 \pm 0.38$ | 0.53 |
| | FixNum | $2.85 \pm 0.82$ | $2.89 \pm 0.86$ | 0.77 |
| | Mean FixDur | $0.20 \pm 0.09$ | $0.18 \pm 0.09$ | 0.29 |
| EEG | Delta | $1.76 \pm 1.23$ | $1.75 \pm 1.22$ | 0.79 |
| | Theta | $18.0 \pm 23.9$ | $15.8 \pm 16.2$ | 0.03 |
| | Alpha | $0.10 \pm 0.08$ | $0.07 \pm 0.07$ | $5.27 \times 10^{-3}$ |
| | Beta | $(5.15 \pm 4.07) \times 10^{-3}$ | $(4.82 \pm 4.02) \times 10^{-3}$ | 0.12 |
| | Gamma | $(1.99 \pm 3.35) \times 10^{-6}$ | $(1.77 \pm 3.13) \times 10^{-6}$ | 0.20 |
| fNIRS | Mean | $0.39 \pm 0.06$ | $0.40 \pm 0.06$ | 0.58 |
| | Variance | $0.08 \pm 0.03$ | $0.08 \pm 0.03$ | 0.48 |
| Skin Conductance | Mean Skin Conductance | $5.30 \pm 2.72$ | $5.79 \pm 2.86$ | 0.24 |
| Respiration | $BR_{coeff}$ | $21.5 \pm 15.4$ | $19.6 \pm 13.2$ | 0.35 |
| | BR | $18.4 \pm 3.01$ | $19.3 \pm 3.42$ | 0.05 |
| | $DC_{coeff}$ | $22.9 \pm 14.6$ | $22.6 \pm 13.4$ | 0.91 |
| | DC | $0.45 \pm 0.14$ | $0.47 \pm 0.14$ | 0.25 |
| | ET | $3.36 \pm 1.92$ | $3.40 \pm 1.69$ | 0.89 |
| | IBI | $3.53 \pm 0.78$ | $3.33 \pm 0.71$ | 0.07 |
| | IE | $0.55 \pm 0.15$ | $0.53 \pm 0.16$ | 0.39 |
| | IT | $1.57 \pm 0.65$ | $1.54 \pm 0.53$ | 0.69 |

(b) Full ANOVA model results.
| Signal Type | Feature | f-value | p-value | Partial Eta Squared |
| --- | --- | --- | --- | --- |
| Pupillometry | APCPS | 81.72 | $2.30 \times 10^{-16}$ | 0.31 |
| | BlinkNum | 0.29 | 0.59 | $1.61 \times 10^{-3}$ |
| | Mean BlinkDur | 0.39 | 0.53 | $2.13 \times 10^{-3}$ |
| | FixNum | 0.08 | 0.77 | $4.52 \times 10^{-4}$ |
| | Mean FixDur | 1.12 | 0.28 | $6.14 \times 10^{-3}$ |
| EEG | Delta | 0.07 | 0.79 | $4.62 \times 10^{-5}$ |
| | Theta | 4.49 | 0.03 | $3.05 \times 10^{-3}$ |
| | Alpha | 7.81 | $5.27 \times 10^{-3}$ | $5.28 \times 10^{-3}$ |
| | Beta | 2.44 | 0.12 | $1.64 \times 10^{-3}$ |
| | Gamma | 1.61 | 0.20 | $1.09 \times 10^{-3}$ |
| fNIRS | Mean | 0.31 | 0.58 | $1.72 \times 10^{-3}$ |
| | Variance | 0.50 | 0.48 | $2.75 \times 10^{-3}$ |
| Skin Conductance | Mean Skin Conductance | 1.42 | 0.23 | $7.75 \times 10^{-3}$ |
| Respiration | $BR_{coeff}$ | 1.30 | 0.26 | $3.01 \times 10^{-3}$ |
|  | BR | 1.65 | 0.20 | 0.01 |
| | $DC_{coeff}$ | 0.08 | 0.78 | $4.46 \times 10^{-4}$ |
|  | DC | 2.03 | 0.16 | 0.01 |
| | ET | 13.99 | $2.52 \times 10^{-4}$ | 0.02 |
| | IBI | 18.91 | $2.43 \times 10^{-3}$ | 0.11 |
| | IE | 16.05 | $9.51 \times 10^{-5}$ | 0.09 |
| | IT | 16.26 | $8.49 \times 10^{-5}$ | 0.09 |

### Workload vs Sense of Urgency

Figure S10 Fig depicts the variation in PCPS taken from the session 1 of a random subject during l_w1_. In the figure, B and D represent the onset of the braking event and the onset of the dialogue event, respectively. We chose l_w1_ to analyze the relationship between the sense of urgency and the workload considering that during l_w1_ of session 1, the participants encountered the braking and the dialogue events for the first time which accounts for a potential increase in the sense of urgency.

**Fig 10.**
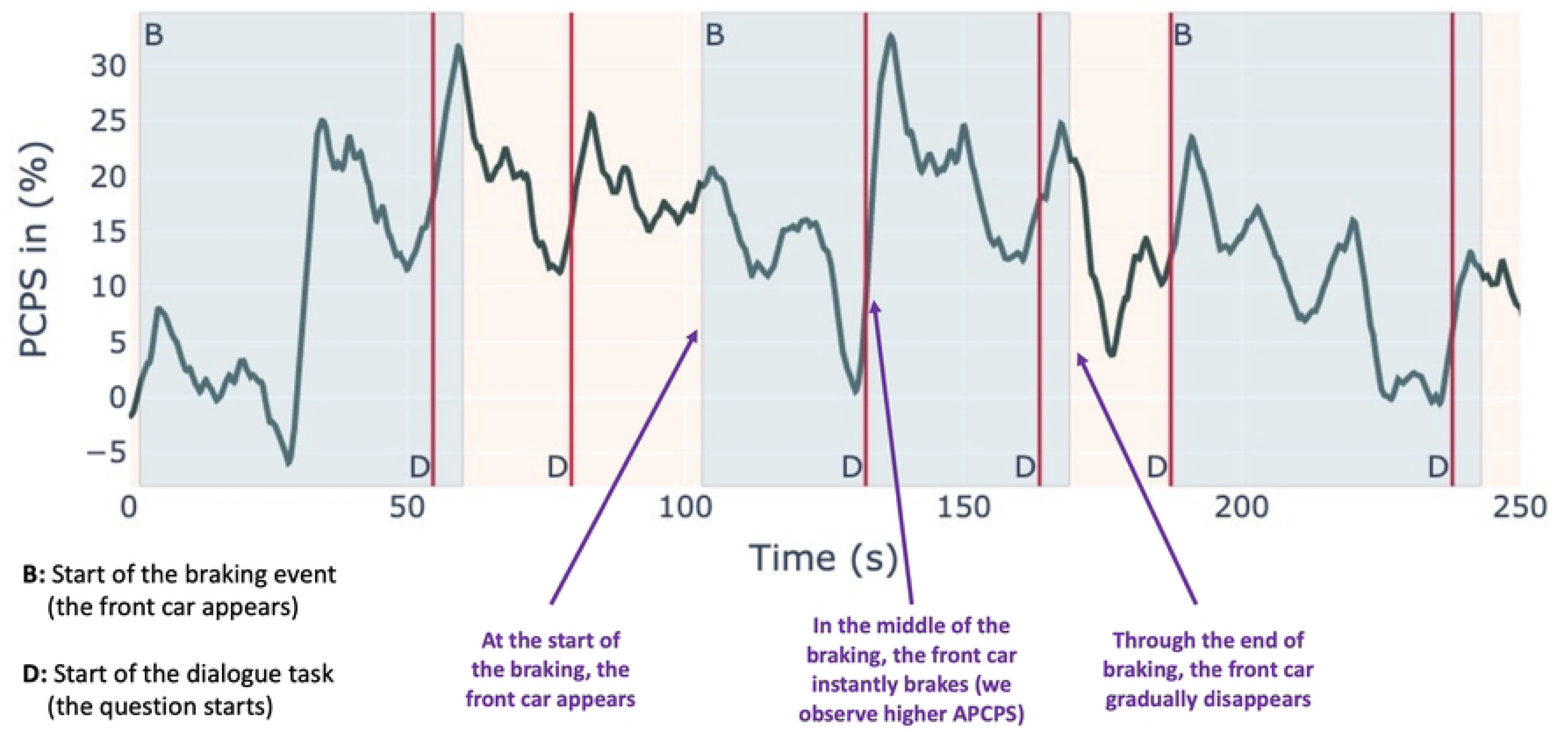
Illustration of the change in PCPS with different events. Illustration of the change in PCPS with different events including braking and dialogue. The results include two sessions obtained from two different subjects. B and D represent the onset of the braking event and the onset of the dialogue event, respectively.

The results indicate that there were noticeable spikes after the onset points of the braking tasks and the dialogue interactions. We observed instantaneous spikes of PCPS in the middle of the braking events. The front car brakes in the middle of the braking event which explains the rapid increase in PCPS during the first braking event of session 1. We also observed instant peaks in PCPS during the second braking event thereupon the dialogue event occurs. The consecutive occurrences of multiple sense of urgency cases thus contribute to higher workload resulting in increased PCPS values during l_w1_ of session 1.

## Model Development and Evaluation

Given that the statistical analyses showed statistically significant differences between the different levels of the three systemic cognitive states, the question arises whether it is possible to train machine learning models on a subset of the data and use the models to predict those levels in new data from new subjects that was not use for training. Put differently, the question is whether state-of-the-art machine-learning methodologies allow for the general prediction of different levels of the three systemic cognitive states across subjects based on the five types of physiological signals: pupillometry, EEG, fNIRS, skin conductance, respiration, and possibly combinations thereof (*e.g.*, pupillometry and EEG). We considered five commonly used models: (1) *k*-Nearest Neighbor (k-NN), (2) Naive-Bayes (NB), (3) Random Forest (RF), (4) Support Vector Machines (SVM), and (5) Multiple Layer Perceptron (MLP) to determine the three human cognitive states (workload, sense of urgency, and mind-wandering) in four cases as follows (described in detail in Table 9):

1. **Three-class workload classification:** We performed the tasks W_S1_ *_−_*_1_, W_S2_ *_−_*_1_, and W_Comb*−*1_.
2. **Two-class workload classification:** We performed the tasks W_S1_ *_−_*_2_, W_S2_ *_−_*_2_, and W_Comb*−*2_.
3. **Two-class sense of urgency classification:** We performed the tasks SoU_S1_, SoU_S2_, and SoU_Comb_.
4. **Two-class mind wandering classification:** We performed the tasks MW_S1_, MW_S2_, and MW_Comb_.

**Table 9.**
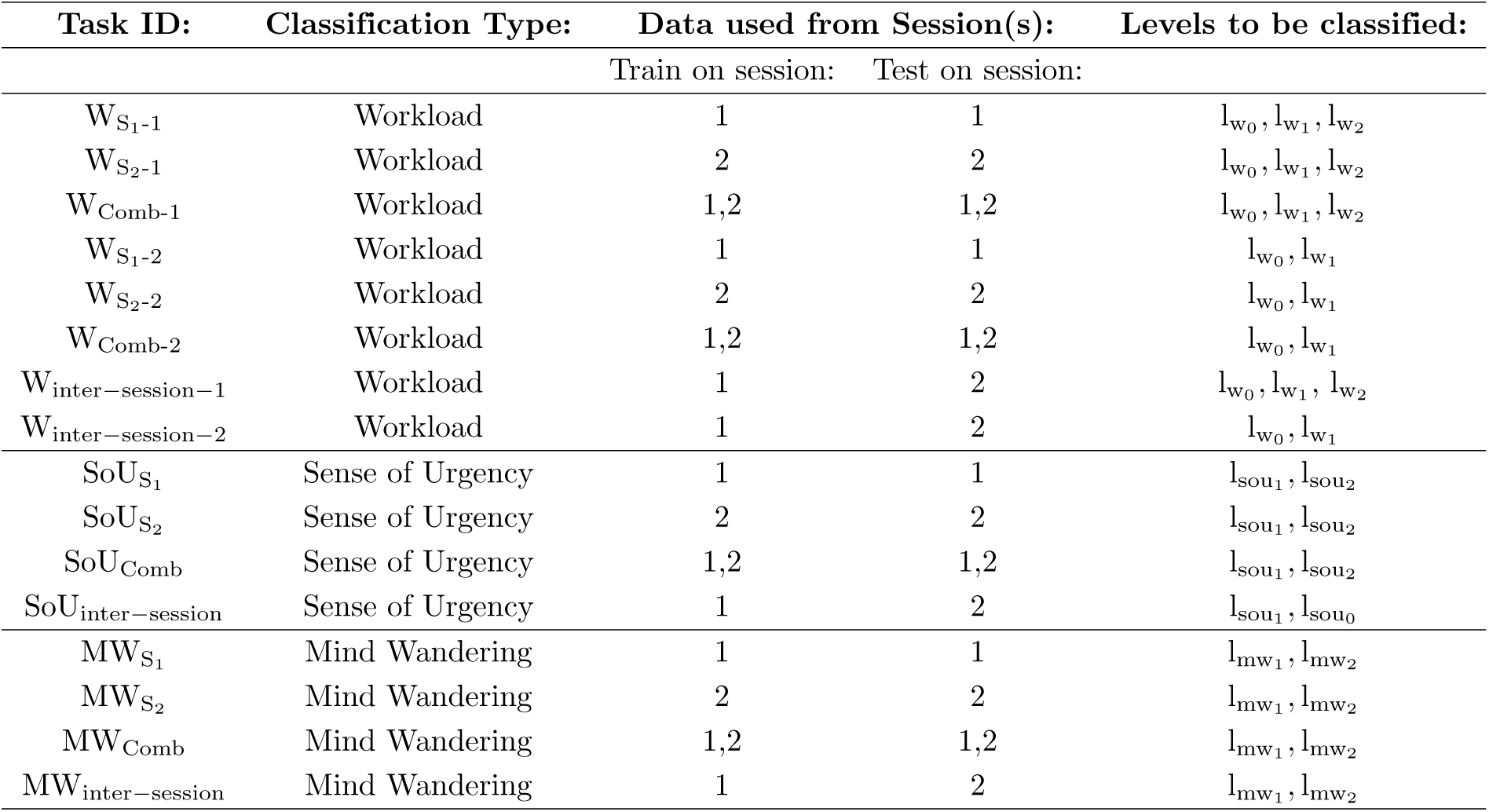
Collection of workload, sense of urgency, and mind wandering classification tasks which includes the task ID, the classification type, the sessions that we used to perform training and testing for classification, and the levels were classified. X_y-z_ represents the classification task of “X” (workload, sense of urgency, or mind wandering) using data from session(s) y (only session 1 (S_1_), only session 2 (S_2_), or the combination of them (Comb) which classifies different levels (1 and 2 represent the three-class classification and the two-class classification of workload, respectively; for mind wandering and sense of urgency, there are only two classes for all classification tasks). As an example, W_S1 -1_ represents the classification of three workload levels obtained from session 1.

For the given classification task, the learning method, and the type of signal, we divided the data into a training set and a test set with a ratio of about 80% and 20%, *i.e.*, data from 37 subjects for the training set and data from 9 subjects for the test set, respectively. Next, the cross-validation method [85] was applied to the training set to select the best-learned model to apply to the test set. To mitigate the randomness in the training and test processes as well as to stabilize the testing performance, for a given classification task and learning method, we repeated the whole experiment 10 times and only report the average accuracy together with its standard deviation. First, we ran each of these tasks separately for 16 features acquired from five signal types which we described in detail in Section “Cognitive State Classification with One Signal Type” under “Development and Evaluation”. Then, in section “Cognitive State Classification with the Fusion of Pupillometry and EEG”, we combined pupillometry and EEG signals and reported the performance of the combination of pupillometry and EEG on twelve classification tasks described in section “Model Development and Evaluation”.

### Cognitive State Classification with One Signal Type

As previously described in the “Methods” Section, we generated three cognitive workload levels (l_w1_, l_w2_, and l_w0_), two sense of urgency levels (l_sou1_ and l_sou2_), and two mind wandering levels (l_mw1_ and l_mw2_) corresponding to different intervals of time and braking/dialogue events (see Figure S3 Fig). Then, we run six workload, three sense of urgency, and three mind wandering classification tasks which we next describe in detail.

#### Three-level Workload Classification

We investigated the performance of the five ML algorithms in separating three cognitive workload levels by using various physiological markers. Table 10a, Table 10b, and Table 10c show the classification accuracies of the five algorithms related to the tasks W_S1_ *_−_*_1_, W_S2_ *_−_*_1_, and W_Comb*−*1_, respectively. The results indicate that for the task W_S1_ *_−_*_1_, which aims to distinguish three workload levels generated from session 1, APCPS achieves the highest accuracy of 72.57 ∓ 8.75 using the Naive Bayes algorithm which outperforms other tested signals by a large margin of nearly 25%. Similarly, for the task W_S2_ *_−_*_1_, which aims to distinguish three workload levels created from session 2, APCPS also achieves the highest accuracy of 59.13 ∓ 8.27, exceeding the accuracy obtained by other signals by a margin of at least nearly 15%. Note that both W_S1_ *_−_*_1_ and W_S2_ *_−_*_1_ are three-class classification tasks, therefore, a random guess will likely provide an accuracy of 33% which is far below the performance achieved by all five tested algorithms on APCPS. Lastly, we run the five ML algorithms for the task W_Comb*−*1_, which aims to distinguish three workload levels by using data from both session 1 and session 2, and found out that APCPS has the best classification accuracy among the 16 features in classifying three workload levels with the highest accuracy of 63.35 ∓ 5.63 using the Naive Bayes algorithm. The results demonstrate that the APCPS signal is the best physiological indicator in separating three cognitive workload levels.

**Table 10.**
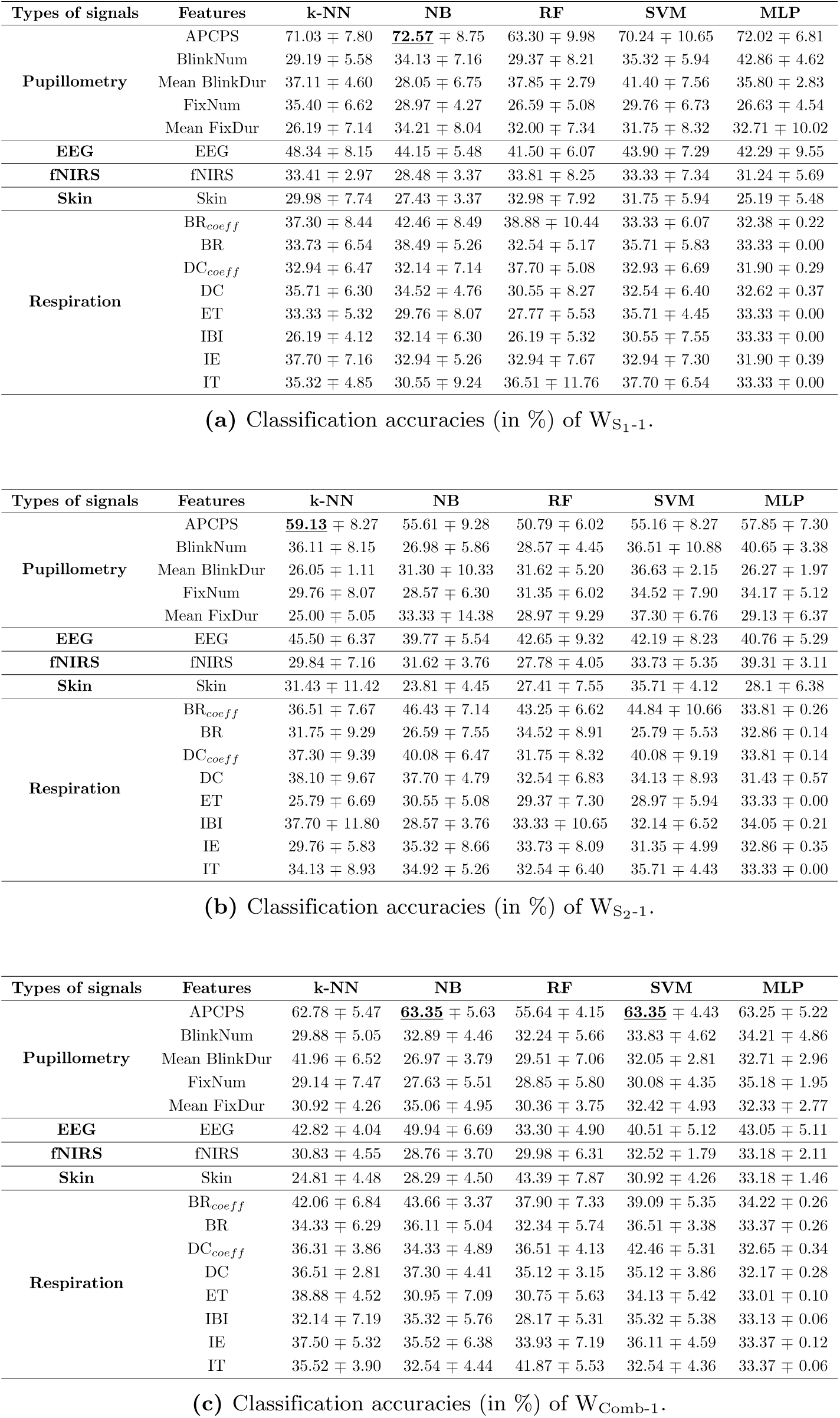
Classification accuracies (in %) of the tasks **(a)** W_S1 -1_, **(b)** W_S2 -1_, and **(c)** W_Comb-1_ described in Table 9.

#### Two-level Workload Classification

Second, we examined the effectiveness of the five ML algorithms in differentiating two cognitive workload levels, l_w1_ and l_w0_, by using different physiological markers. Table 11a, Table 11b, and Table 11c show the classification accuracies of five algorithms related to the tasks W_S1_ *_−_*_2_, W_S2_ *_−_*_2_, and W_Comb*−*2_, respectively. For the classification tasks W_S1_ *_−_*_2_ and W_S2_ *_−_*_2_, that aim to discern between the lowest and the highest workload levels in session 1 and session 2, respectively, the APCPS outperforms other signals by a large margin and achieves impressive accuracies of 93.25 ∓ 3.55 (W_S1_ *_−_*_2_) by using MLP and 88.89 ∓ 5.23 (W_S2_ *_−_*_2_) by using NB. Similarly, the APCPS achieved the best classification accuracy (90.18 ∓ 3.42) for the task W_Comb*−*2_ which aims to separate l_w1_ and l_w0_ by using the data obtained from both session 1 and session 2. The observed numerical results in tasks W_S1_ *_−_*_2_, W_S2_ *_−_*_2_, and W_Comb*−*2_ consistently demonstrate the superior performance of APCPS in workload classification tasks compared to other tested physiological signals.

**Table 11.** Classification accuracies (in %) of the tasks **(a)** W_S1 -2_, **(b)** W_S2 -2_, and **(c)** W_Comb-2_ described in Table 9.

| Types of signals | Features | k-NN | NB | RF | SVM | MLP |
| --- | --- | --- | --- | --- | --- | --- |
| **Pupillometry** | APCPS | 88.89 ± 6.77 | 90.06 ± 5.23 | 89.47 ± 6.56 | 92.98 ± 4.96 | **93.25** ± 3.55 |
| | BlinkNum | 53.86 ± 6.08 | 53.22 ± 5.79 | 49.77 ± 7.49 | 50.29 ± 8.63 | 50.40 ± 6.17 |
| | Mean BlinkDur | 55.72 ± 12.10 | 49.27 ± 4.70 | 59.12 ± 10.22 | 52.19 ± 12.23 | 56.17 ± 9.00 |
| | FixNum | 52.05 ± 8.02 | 47.37 ± 7.44 | 48.54 ± 7.35 | 42.11 ± 8.59 | 50.00 ± 0.00 |
| | Mean FixDur | 53.86 ± 7.85 | 43.86 ± 6.56 | 43.86 ± 14.25 | 42.69 ± 9.10 | 51.19 ± 6.52 |
| **EEG** | EEG | 61.27 ± 4.33 | 58.60 ± 5.51 | 57.46 ± 3.83 | 60.98 ± 7.35 | 60.18 ± 5.52 |
| **fNIRS** | fNIRS | 49.77 ± 7.07 | 59.49 ± 3.51 | 55.32 ± 4.60 | 50.29 ± 10.85 | 52.05 ± 8.02 |
| **Skin** | Skin | 46.26 ± 11.22 | 45.03 ± 5.03 | 45.03 ± 13.84 | 44.44 ± 8.27 | 49.21 ± 1.48 |
| **Respiration** | BR<sub>coeff</sub> | 56.21 ± 9.16 | 54.39 ± 10.81 | 60.82 ± 9.64 | 64.91 ± 9.92 | 47.37 ± 7.44 |
| | BR | 47.37 ± 10.81 | 45.03 ± 9.64 | 40.94 ± 9.21 | 43.86 ± 7.85 | 48.95 ± 2.41 |
| | DC<sub>coeff</sub> | 53.80 ± 14.42 | 48.54 ± 8.51 | 60.23 ± 10.26 | 56.14 ± 8.59 | 48.42 ± 3.16 |
| | DC | 59.06 ± 8.51 | 56.14 ± 9.28 | 49.12 ± 11.10 | 50.29 ± 8.27 | 48.42 ± 5.16 |
| | ET | 57.31 ± 8.02 | 46.78 ± 5.79 | 47.95 ± 7.62 | 57.31 ± 7.21 | 50.53 ± 2.58 |
| | IBI | 46.20 ± 6.46 | 45.61 ± 13.36 | 36.26 ± 10.36 | 42.69 ± 6.30 | 50.00 ± 2.63 |
| | IE | 47.95 ± 7.21 | 57.31 ± 10.06 | 59.06 ± 12.35 | 52.05 ± 12.27 | 46.32 ± 12.85 |
| | IT | 59.06 ± 8.51 | 48.54 ± 8.87 | 47.37 ± 11.10 | 52.05 ± 7.62 | 50.53 ± 3.49 |

(a) Classification accuracies (in %) of $W_{S_1-2}$ .
| Types of signals | Features | k-NN | NB | RF | SVM | MLP |
| --- | --- | --- | --- | --- | --- | --- |
| **Pupillometry** | APCPS | 86.55 ± 5.61 | **88.89** ± 5.23 | 84.80 ± 6.30 | 84.80 ± 4.60 | 84.52 ± 4.45 |
| | BlinkNum | 37.43 ± 7.21 | 46.20 ± 5.96 | 46.78 ± 6.30 | 46.20 ± 7.35 | 46.03 ± 6.17 |
| | Mean BlinkDur | 48.12 ± 7.37 | 52.86 ± 3.33 | 61.10 ± 5.55 | 46.60 ± 6.85 | 49.52 ± 2.38 |
| | FixNum | 48.54 ± 9.86 | 40.94 ± 8.15 | 46.78 ± 6.30 | 45.03 ± 8.27 | 50.00 ± 1.68 |
| | Mean FixDur | 48.01 ± 6.46 | 42.11 ± 8.95 | 47.95 ± 8.02 | 43.27 ± 8.87 | 48.41 ± 5.08 |
| **EEG** | EEG | 56.25 ± 9.19 | 52.30 ± 4.81 | 52.60 ± 5.08 | 58.15 ± 2.65 | 54.33 ± 3.17 |
| **fNIRS** | fNIRS | 42.16 ± 9.10 | 56.42 ± 1.65 | 58.83 ± 2.19 | 47.37 ± 3.51 | 53.00 ± 4.01 |
| **Skin** | Skin | 44.01 ± 7.35 | 43.27 ± 8.51 | 45.61 ± 15.49 | 42.11 ± 8.23 | 48.81 ± 3.37 |
| **Respiration** | BR<sub>coeff</sub> | 56.14 ± 9.92 | 61.99 ± 11.03 | 55.55 ± 8.63 | 50.88 ± 9.61 | 50.53 ± 3.49 |
| | BR | 49.12 ± 9.92 | 44.44 ± 12.18 | 41.52 ± 10.66 | 37.43 ± 9.10 | 51.05 ± 4.74 |
| | DC<sub>coeff</sub> | 46.78 ± 9.10 | 52.05 ± 9.10 | 45.61 ± 10.23 | 52.05 ± 8.02 | 48.95 ± 3.27 |
| | DC | 47.95 ± 9.75 | 40.94 ± 7.76 | 53.22 ± 12.76 | 39.77 ± 8.63 | 46.84 ± 5.49 |
| | ET | 39.18 ± 9.32 | 46.20 ± 7.76 | 32.75 ± 5.42 | 45.61 ± 3.51 | 50.53 ± 2.58 |
| | IBI | 43.27 ± 7.76 | 42.69 ± 5.23 | 50.29 ± 10.26 | 46.20 ± 9.86 | 51.58 ± 6.14 |
| | IE | 42.69 ± 6.77 | 43.86 ± 12.89 | 47.95 ± 9.43 | 45.61 ± 7.85 | 46.84 ± 5.50 |
| | IT | 38.01 ± 12.10 | 46.78 ± 4.60 | 42.11 ± 9.61 | 42.69 ± 4.60 | 48.95 ± 2.41 |

(b) Classification accuracies (in %) of $W_{S_2-2}$ .
| Types of signals | Features | k-NN | NB | RF | SVM | MLP |
| --- | --- | --- | --- | --- | --- | --- |
| **Pupillometry** | APCPS | **90.18** ± 3.42 | 87.91 ± 6.04 | 85.35 ± 5.97 | 88.76 ± 4.14 | 85.71 ± 4.41 |
| | BlinkNum | 49.22 ± 7.47 | 51.64 ± 7.59 | 49.79 ± 7.09 | 50.92 ± 7.71 | 51.03 ± 4.03 |
| | Mean BlinkDur | 61.09 ± 4.36 | 45.80 ± 5.15 | 44.10 ± 6.62 | 47.65 ± 4.82 | 48.68 ± 2.77 |
| | FixNum | 46.80 ± 5.62 | 43.39 ± 6.23 | 49.22 ± 7.37 | 49.36 ± 5.94 | 49.99 ± 1.00 |
| | Mean FixDur | 44.52 ± 7.04 | 43.53 ± 7.69 | 48.22 ± 7.40 | 48.93 ± 6.44 | 49.99 ± 5.50 |
| **EEG** | EEG | 60.95 ± 5.41 | 58.95 ± 5.83 | 54.49 ± 7.71 | 63.44 ± 5.64 | 63.44 ± 4.19 |
| **fNIRS** | fNIRS | 47.52 ± 5.56 | 45.09 ± 5.95 | 42.25 ± 7.08 | 49.34 ± 1.44 | 49.81 ± 2.45 |
| **Skin** | Skin | 40.26 ± 6.73 | 44.81 ± 4.32 | 28.67 ± 6.20 | 48.36 ± 6.84 | 49.25 ± 3.44 |
| **Respiration** | BR<sub>coeff</sub> | 55.26 ± 5.57 | 50.15 ± 5.11 | 56.76 ± 9.62 | 51.35 ± 7.09 | 48.11 ± 3.38 |
| | BR | 48.95 ± 3.92 | 47.45 ± 4.61 | 44.14 ± 8.16 | 48.65 ± 8.26 | 51.08 ± 3.07 |
| | DC<sub>coeff</sub> | 53.45 ± 5.94 | 50.45 ± 5.70 | 46.55 ± 5.95 | 48.05 ± 6.34 | 49.46 ± 2.72 |
| | DC | 55.55 ± 8.56 | 54.65 ± 5.22 | 48.05 ± 5.95 | 51.65 ± 9.40 | 49.46 ± 6.40 |
| | ET | 50.75 ± 5.06 | 49.55 ± 7.10 | 51.05 ± 8.96 | 54.05 ± 7.54 | 48.19 ± 2.65 |
| | IBI | 47.45 ± 8.46 | 42.94 ± 6.17 | 41.74 ± 6.51 | 48.65 ± 5.41 | 49.19 ± 1.08 |
| | IE | 52.25 ± 6.98 | 52.85 ± 5.42 | 59.46 ± 5.41 | 53.45 ± 9.26 | 48.65 ± 6.28 |
| | IT | 49.25 ± 4.90 | 50.15 ± 3.63 | 55.26 ± 5.27 | 52.85 ± 5.27 | 50.27 ± 1.32 |

(c) Classification accuracies (in %) of $W_{Comb-2}$ .

#### Two-level Sense of Urgency Classification

We defined two different levels for sense of urgency for use with the five ML algorithms. Table 12a, Table 12b, and Table 12c provide the classification accuracies of five algorithms related to the tasks SoU_S1_, SoU_S2_, and SoU_Comb_, respectively. Specifically, the highest classification accuracy for task SoU_S1_, task SoU_S2_, and task SoU_Comb_ are 63.43%, 64.27%, and 60.88% achieved by FixNum, APCPS and Mean FixDur, respectively. Even though the classification performance may depend on the learning algorithm, it seems like APCSP and Mean FixDur are the best two signals for assessing the sense of urgency that not only provide higher accuracy but also more stability than other compared signals.

**Table 12.** Classification accuracies (in %) of the tasks **(a)**SoU_S1_, **(b)** SoU_S2_, and **(c)** SoU_Comb_ described in Table 9.

| Types of signals | Features | k-NN | NB | RF | SVM | MLP |
| --- | --- | --- | --- | --- | --- | --- |
| Pupillometry | APCPS | 51.08 ± 12.65 | 55.96 ± 7.48 | 44.32 ± 9.09 | 49.86 ± 9.56 | 48.42 ± 10.00 |
| | FixNum | 50.80 ± 10.56 | 61.77 ± 9.33 | 52.91 ± 12.37 | 63.43 ± 9.58 | 52.57 ± 9.39 |
| | Mean FixDur | 61.88 ± 15.13 | 59.83 ± 13.68 | 52.63 ± 10.10 | 59.56 ± 9.20 | 59.06 ± 10.42 |
| EEG | EEG | 48.14 ± 7.32 | 51.52 ± 11.90 | 53.02 ± 10.08 | 49.80 ± 7.30 | 48.25 ± 7.41 |
| fNIRS | fNIRS | 44.75 ± 11.43 | 49.46 ± 8.06 | 48.63 ± 8.19 | 45.42 ± 6.90 | 43.14 ± 5.50 |
| Skin | Skin | 32.90 ± 10.69 | 44.60 ± 7.32 | 44.32 ± 9.09 | 50.42 ± 7.88 | 48.14 ± 5.76 |

(a) Classification accuracies (in %) of SoU<sub>S<sub>1</sub></sub>.
| Types of signals | Features | k-NN | NB | RF | SVM | MLP |
| --- | --- | --- | --- | --- | --- | --- |
| Pupillometry | APCPS | 63.82 ± 7.05 | 64.27 ± 10.88 | 55.40 ± 10.01 | 63.16 ± 8.87 | 58.39 ± 7.71 |
| | FixNum | 59.66 ± 9.41 | 49.03 ± 9.96 | 60.94 ± 9.71 | 44.88 ± 8.08 | 45.65 ± 8.95 |
| | Mean FixDur | 56.62 ± 5.78 | 56.23 ± 11.20 | 51.52 ± 11.01 | 54.29 ± 7.65 | 53.06 ± 9.70 |
| EEG | EEG | 56.73 ± 9.55 | 59.83 ± 10.28 | 57.89 ± 13.00 | 58.39 ± 9.25 | 57.11 ± 11.50 |
| fNIRS | fNIRS | 56.94 ± 9.09 | 57.31 ± 10.47 | 52.69 ± 9.84 | 54.56 ± 9.96 | 58.10 ± 8.05 |
| Skin | Skin | 42.49 ± 8.34 | 43.49 ± 5.08 | 46.81 ± 8.69 | 45.98 ± 9.48 | 46.20 ± 4.60 |

(b) Classification accuracies (in %) of SoU<sub>S<sub>2</sub></sub>.
| Types of signals | Features | k-NN | NB | RF | SVM | MLP |
| --- | --- | --- | --- | --- | --- | --- |
| Pupillometry | APCPS | 57.04 ± 6.52 | 60.46 ± 9.29 | 54.05 ± 5.95 | 57.04 ± 8.41 | 55.75 ± 7.80 |
| | FixNum | 55.90 ± 7.62 | 53.91 ± 5.51 | 57.33 ± 7.15 | 55.33 ± 8.87 | 50.49 ± 4.14 |
| | Mean FixDur | 56.61 ± 8.77 | 59.17 ± 6.32 | 52.63 ± 7.09 | 60.88 ± 8.34 | 58.17 ± 8.17 |
| EEG | EEG | 54.35 ± 5.61 | 59.60 ± 9.74 | 51.06 ± 7.88 | 51.34 ± 5.03 | 56.74 ± 6.67 |
| fNIRS | fNIRS | 50.92 ± 6.52 | 55.04 ± 7.36 | 45.66 ± 8.45 | 50.35 ± 8.14 | 53.61 ± 9.96 |
| Skin | Skin | 41.11 ± 5.70 | 46.23 ± 7.28 | 47.23 ± 5.64 | 47.51 ± 6.70 | 46.08 ± 4.26 |

(c) Classification accuracies (in %) of SoU<sub>Comb</sub>.

#### Two-level Mind Wandering Classification

Two different levels of mind wandering were defined based on the first 1.5 minutes (l_mw1_) and the second 1.5 minutes (l_mw2_) of the experiment in each session, respectively. We defined three mind wandering classification tasks, MW_S1_, MW_S2_, and MW_Comb_ which aimed to differentiate two mind wandering levels by utilizing the data acquired from session 1, from session 2, and from the combination of session 1 and session 2, respectively. The mind wandering evaluation design is outlined in the Methods section.

We refer the readers to Table 9 for the details of all of the classification tasks. Table 13a, Table 13b, and Table 13c provide the mind wandering classification accuracies corresponding to three classification tasks MW_S1_, MW_S2_, and MW_Comb_, respectively. The results indicate that for the task MW_S1_ which aims to classify between two mind-wandering levels in session 1, the APCPS achieves the highest accuracy of 76.22 ∓ 4.17 (with RF) which outperforms all other tested signals by at least 10%. Similarly, we observed the same situation where the APCPS provides significantly better performance compared to other signals on the classification task MW_S2_. Particularly, the highest classification accuracy between the two mind-wandering levels in session 2 is 86.08 ∓ 5.55 achieved by the APCPS using the RF algorithm. This outperforms other tested signals by a large gap of at least 20%. In addition, the APCPS has the best accuracy (82.08 ∓ 7.35) in differentiating the two mind wandering levels by using the data obtained from both session 1 and session 2. The results of the mind wandering classification tasks demonstrate that the APCPS has the best performance in predicting two mind wandering levels regardless of which classification model is used.

**Table 13.**
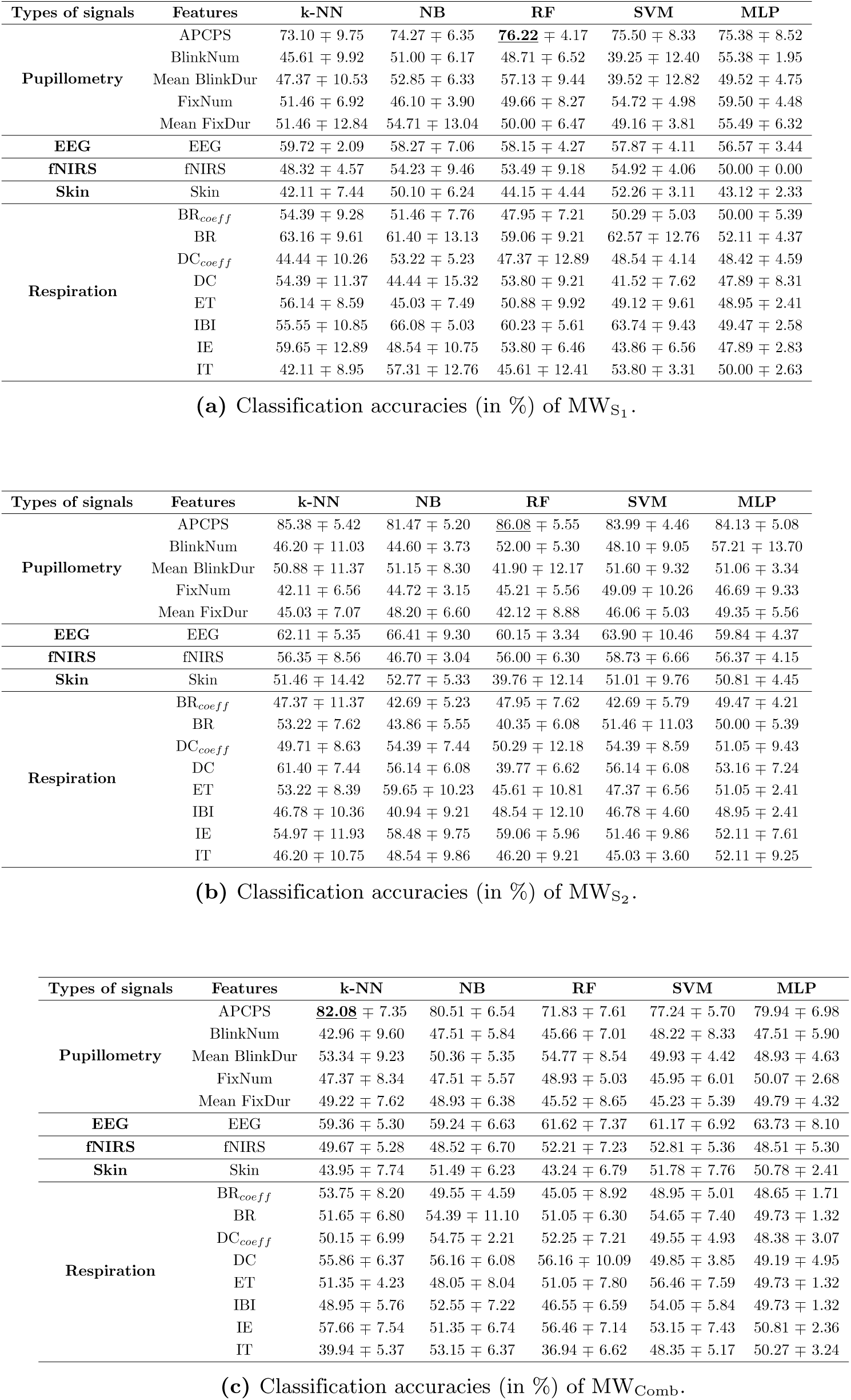
Classification accuracies (in %) of the tasks **(a)** MW_S1_, **(b)** MW_S2_, and **(c)** MW_Comb_ described in Table 9.

### Assessing the Performance of Inter-State Class Classification

We further examined the performance of different ML/DL methods in differentiating multiple levels of cognitive states by training the models on physiological biomarkers from Session 1 and evaluating them using biomarkers from Session 2. In particular, the inter-session classification tasks are defined as follows: W_inter*−*session*−*1_ represents the three-class workload classification, where the ML/DL models are trained using data from Session 1 and evaluated on data from Session 2; W_inter*−*session*−*2_ denotes the two-class workload classification under the same inter-session training and testing scheme. Similarly, SoU_inter*−*session_ corresponds to the two-class sense of urgency classification, and MW_inter*−*session_ refers to the two-class mind wandering classification, both trained on Session 1 and tested on Session 2. The detailed task definitions are summarized in Table 9. The results indicated that PCPS continues to demonstrate superior performance in classifying different levels of the three cognitive states including workload, sense of urgency, and mind wandering see Tables 14a, 14b, 14c, and 14d.

**Table 14.** Classification accuracies (in %) of the tasks **(a)** W_inter*−*session*−*1_, **(b)** W_inter*−*session*−*2_, **(c)** SoU_inter*−*session_, and **(d)** MW_inter*−*session_ described in Table 9.

| Types of signals | Features | k-NN | NB | RF | SVM | MLP |
| --- | --- | --- | --- | --- | --- | --- |
| Pupillometry | APCPS | 83.09 ± 2.06 | 84.06 ± 3.89 | 83.09 ± 1.37 | 84.06 ± 1.51 | 83.82 ± 3.80 |
| | BlinkNum | 53.74 ± 2.06 | 49.99 ± 1.26 | 59.54 ± 1.43 | 49.76 ± 2.04 | 49.26 ± 1.85 |
| | Mean BlinkDur | 41.42 ± 2.93 | 47.04 ± 1.76 | 45.83 ± 5.32 | 39.35 ± 4.62 | 37.51 ± 3.49 |
| | FixNum | 49.03 ± 2.26 | 46.86 ± 1.49 | 50.12 ± 2.87 | 50.12 ± 2.87 | 52.05 ± 2.20 |
| | Mean FixDur | 49.88 ± 3.34 | 49.64 ± 1.77 | 49.64 ± 1.77 | 50.48 ± 2.46 | 49.88 ± 3.14 |
| EEG | EEG | 57.79 ± 2.95 | 47.04 ± 1.82 | 52.13 ± 2.56 | 45.35 ± 3.61 | 43.66 ± 2.88 |
| fNIRS | fNIRS | 49.88 ± 3.88 | 49.88 ± 8.01 | 44.08 ± 1.55 | 50.85 ± 1.43 | 50.72 ± 1.02 |
| Skin | Skin | 47.95 ± 1.88 | 50.0 ± 1.26 | 46.26 ± 4.56 | 49.40 ± 1.37 | 49.64 ± 0.72 |
| Respiration | BR<sub>coeff</sub> | 55.07 ± 1.92 | 63.04 ± 2.89 | 57.73 ± 2.53 | 62.68 ± 1.02 | 62.44 ± 1.46 |
| | BR | 61.71 ± 4.91 | 58.21 ± 3.18 | 56.64 ± 1.81 | 60.63 ± 4.16 | 64.13 ± 3.44 |
| | DC<sub>coeff</sub> | 53.74 ± 1.78 | 59.42 ± 5.40 | 55.80 ± 2.05 | 54.23 ± 5.99 | 57.61 ± 4.07 |
| | DC | 60.51 ± 1.62 | 62.20 ± 1.00 | 56.16 ± 3.44 | 63.53 ± 3.54 | 64.01 ± 0.62 |
| | ET | 54.11 ± 9.69 | 62.44 ± 3.54 | 42.75 ± 4.07 | 61.71 ± 2.68 | 61.96 ± 1.51 |
| | IBI | 56.76 ± 1.76 | 60.87 ± 7.11 | 44.57 ± 4.03 | 60.35 ± 0.72 | 61.00 ± 1.34 |
| | IE | 60.63 ± 0.85 | 60.99 ± 3.42 | 55.43 ± 2.05 | 59.25 ± 3.60 | 53.91 ± 3.05 |
| | IT | 58.70 ± 3.07 | 57.31 ± 3.46 | 53.74 ± 4.26 | 58.82 ± 7.40 | 55.74 ± 2.16 |

(a) Classification accuracies (in %) of $W_{\text{inter-session-1}}$ .
| Types of signals | Features | k-NN | NB | RF | SVM | MLP |
| --- | --- | --- | --- | --- | --- | --- |
| Pupillometry | APCPS | 55.24 ± 4.96 | 59.18 ± 3.97 | 58.29 ± 2.96 | 58.10 ± 2.82 | 60.95 ± 1.69 |
| | BlinkNum | 33.17 ± 5.04 | 35.99 ± 1.23 | 32.69 ± 3.75 | 33.73 ± 6.10 | 37.44 ± 2.27 |
| | Mean BlinkDur | 36.88 ± 5.72 | 33.17 ± 1.80 | 36.88 ± 4.10 | 39.55 ± 1.17 | 41.93 ± 3.71 |
| | FixNum | 33.82 ± 4.16 | 32.05 ± 4.50 | 27.05 ± 4.21 | 29.37 ± 3.27 | 33.01 ± 1.68 |
| | Mean FixDur | 28.99 ± 4.78 | 27.21 ± 2.02 | 28.18 ± 4.33 | 34.92 ± 9.78 | 35.75 ± 1.84 |
| EEG | EEG | 43.07 ± 0.69 | 41.14 ± 2.46 | 40.50 ± 7.34 | 43.00 ± 1.76 | 38.43 ± 5.16 |
| fNIRS | fNIRS | 34.67 ± 3.21 | 41.61 ± 3.24 | 31.77 ± 4.16 | 33.10 ± 3.37 | 36.34 ± 3.60 |
| Skin | Skin | 28.66 ± 3.40 | 30.11 ± 4.36 | 30.76 ± 4.52 | 31.35 ± 2.24 | 32.85 ± 4.08 |
| Respiration | BR<sub>coeff</sub> | 36.23 ± 5.84 | 35.75 ± 3.55 | 33.98 ± 4.22 | 36.51 ± 4.99 | 41.22 ± 1.62 |
| | BR | 37.36 ± 6.79 | 38.33 ± 2.91 | 38.49 ± 4.11 | 38.49 ± 7.49 | 37.60 ± 3.34 |
| | DC<sub>coeff</sub> | 36.55 ± 4.67 | 31.56 ± 4.36 | 38.81 ± 3.47 | 42.54 ± 5.70 | 42.75 ± 4.18 |
| | DC | 42.67 ± 5.80 | 38.49 ± 5.30 | 34.62 ± 3.58 | 43.66 ± 5.53 | 45.81 ± 7.75 |
| | ET | 42.03 ± 5.88 | 43.00 ± 2.65 | 40.90 ± 5.37 | 43.25 ± 6.62 | 42.11 ± 2.89 |
| | IBI | 39.45 ± 7.09 | 44.12 ± 2.38 | 37.52 ± 3.38 | 41.66 ± 6.07 | 40.42 ± 0.46 |
| | IE | 41.38 ± 4.54 | 42.35 ± 3.91 | 39.14 ± 3.48 | 44.84 ± 5.35 | 41.71 ± 1.45 |
| | IT | 41.22 ± 5.25 | 42.67 ± 4.88 | 34.94 ± 5.22 | 39.29 ± 4.45 | 41.47 ± 5.12 |

(b) Classification accuracies (in %) of $W_{\text{inter-session-2}}$ .
| Types of signals | Features | k-NN | NB | RF | SVM | MLP |
| --- | --- | --- | --- | --- | --- | --- |
| Pupillometry | APCPS | 55.63 ± 4.97 | 58.81 ± 2.81 | 51.63 ± 4.59 | 58.92 ± 4.20 | 51.12 ± 5.50 |
| | FixNum | 57.18 ± 5.13 | 53.60 ± 5.33 | 53.75 ± 4.38 | 59.95 ± 3.80 | 45.29 ± 3.13 |
| | Mean FixDur | 59.81 ± 4.24 | 59.55 ± 3.97 | 50.89 ± 3.84 | 61.55 ± 4.21 | 55.93 ± 6.45 |
| EEG | EEG | 59.64 ± 6.22 | 58.58 ± 4.96 | 50.95 ± 4.66 | 58.30 ± 5.93 | 57.47 ± 2.72 |
| fNIRS | fNIRS | 60.38 ± 6.36 | 58.52 ± 7.96 | 51.46 ± 3.92 | 58.27 ± 6.17 | 54.70 ± 3.53 |
| Skin | Skin | 45.62 ± 3.87 | 48.85 ± 3.59 | 46.37 ± 5.51 | 54.68 ± 3.69 | 45.23 ± 1.60 |

(c) Classification accuracies (in %) of $SoU_{\text{inter-session}}$ .
| Types of signals | Features | k-NN | NB | RF | SVM | MLP |
| --- | --- | --- | --- | --- | --- | --- |
| Pupillometry | APCPS | 72.03 ± 3.71 | 74.61 ± 0.72 | 76.77 ± 2.78 | 78.09 ± 1.04 | 74.45 ± 1.52 |
| | BlinkNum | 45.17 ± 3.92 | 47.83 ± 2.81 | 46.01 ± 3.87 | 49.88 ± 3.05 | 49.52 ± 2.67 |
| | Mean BlinkDur | 39.72 ± 3.93 | 52.48 ± 1.77 | 48.50 ± 2.14 | 55.22 ± 2.33 | 41.11 ± 3.19 |
| | FixNum | 51.81 ± 1.62 | 49.64 ± 2.61 | 49.64 ± 2.94 | 47.58 ± 3.89 | 51.93 ± 4.06 |
| | Mean FixDur | 50.92 ± 4.80 | 46.66 ± 3.16 | 47.95 ± 2.53 | 50.72 ± 1.70 | 46.01 ± 1.15 |
| EEG | EEG | 55.10 ± 1.75 | 44.38 ± 2.63 | 40.27 ± 2.21 | 47.36 ± 1.09 | 53.01 ± 2.68 |
| fNIRS | fNIRS | 53.86 ± 1.74 | 36.46 ± 1.24 | 38.97 ± 2.13 | 46.07 ± 3.05 | 45.75 ± 1.87 |
| Skin | Skin | 51.33 ± 3.19 | 50.97 ± 2.34 | 49.03 ± 2.08 | 51.93 ± 1.00 | 53.16 ± 1.89 |
| Respiration | BR<sub>coeff</sub> | 46.08 ± 2.76 | 49.76 ± 1.33 | 49.88 ± 3.56 | 50.00 ± 2.94 | 53.72 ± 3.03 |
| | BR | 47.11 ± 3.92 | 49.40 ± 1.04 | 47.71 ± 4.30 | 49.76 ± 1.33 | 49.28 ± 2.17 |
| | DC<sub>coeff</sub> | 46.13 ± 1.86 | 45.53 ± 4.34 | 51.09 ± 2.40 | 49.28 ± 5.07 | 52.48 ± 4.35 |
| | DC | 53.86 ± 0.74 | 54.35 ± 0.0 | 55.19 ± 1.23 | 52.66 ± 2.18 | 53.74 ± 0.90 |
| | ET | 55.19 ± 3.15 | 57.13 ± 5.54 | 50.60 ± 3.84 | 57.37 ± 0.68 | 56.00 ± 1.78 |
| | IBI | 57.73 ± 2.26 | 59.06 ± 1.02 | 57.25 ± 1.98 | 59.90 ± 1.08 | 58.66 ± 1.20 |
| | IE | 58.70 ± 4.10 | 57.94 ± 6.02 | 51.57 ± 2.18 | 58.94 ± 1.90 | 60.14 ± 1.85 |
| | IT | 56.52 ± 1.91 | 58.95 ± 3.71 | 55.92 ± 2.72 | 58.82 ± 0.84 | 58.57 ± 0.34 |

(d) Classification accuracies (in %) of $MW_{\text{inter-session-2}}$ .

### Cognitive State Classification with the Fusion of Pupillometry and EEG

We also investigated whether combining different types of physiological signal modalities could improve the performance of single modalities in workload, sense of urgency, and mind wandering classification tasks. Since we had 16 physiological signals, there were numerous combinations which may potentially result in a very high computational complexity. Therefore, we limited ourselves to only combining the APCPS and EEG given that they were the best two indicators for assessing different human cognitive state levels. I.e., even though respiration features including BR_coeff_, BR, DC_coeff_, and DC had slightly better accuracies compared to EEG, considering all of the twelve classification tasks (described in section “Development and Evaluation”), EEG is still the second best physiological indicator which guided us to combine the APCPS and EEG. Next, we combined the APCPS and EEG signals given in section “Physiological Signal Modalities” to form a single input and performed all five ML algorithms (k-NN, NB, RF, SVM, MLP) for all six workload, three sense of urgency, and three mind wandering classification tasks (described in Table 9). The results are shown in Table 15. For all workload tasks (W_S1 -1_, W_S2 -1_, W_Comb-1_, W_S1 -2_, W_S2 -2_, and W_Comb-2_) and mind wandering tasks (MW_S1_, MW_S2_, and MW_Comb_), combining the APCPS and EEG did not lead to better performance compared to APCPS alone. A similar phenomenon can be observed in a sense of urgency when combining APCPS and Mean FixDur which did not significantly improve the overall classification performance. For example, the highest accuracy for the APCPS in task W_S1 -2_ is 93.25 ∓ 3.55 (see Table 11a). However, the highest accuracy in task W_S1 -2_ by combining the APCPS and EEG is just 89.20 ∓ 4.62 (see Table 15) which is 4% lower. However, it is worth noting that the performance of this combination largely outperforms the performance obtained by EEG only. For example, the highest accuracy for EEG in task W_S2 -2_ (see Table 11b) is just 58.15 ∓ 2.65, however, the highest accuracy in task W_S2 -2_ by combining the APCPS and EEG is 85.44 ∓ 5.48 (see Table 15). Again, this shows that APCPS, and not EEG, is really the important modality responsibility for the high classification performance.

**Table 15.** Classification accuracies (in %) by combining APCPS and other physiological signals (EEG, Mean FixDur) for six workload, three sense of urgency, and three mind wandering classification tasks defined in Table 9 by different ML methods.

| Signal Modality | Tasks | k-NN | NB | RF | SVM | MLP |
| --- | --- | --- | --- | --- | --- | --- |
| Workload (APCPS+EEG) | W<sub>S1-1</sub> | 64.56 ± 8.02 | 66.50 ± 4.54 | 65.15 ± 8.33 | 67.12 ± 3.22 | 67.30 ± 7.36 |
| | W<sub>S2-1</sub> | 57.92 ± 5.80 | 56.03 ± 1.72 | 52.44 ± 4.67 | 54.81 ± 5.52 | 57.80 ± 6.27 |
| | W<sub>Comb-1</sub> | 61.03 ± 5.93 | 60.97 ± 4.63 | 52.04 ± 6.34 | 63.26 ± 5.63 | 62.38 ± 4.16 |
| Workload (APCPS+EEG) | W<sub>S1-2</sub> | 86.55 ± 10.10 | 88.63 ± 9.21 | 89.41 ± 4.19 | 85.97 ± 6.35 | 89.20 ± 4.62 |
| | W<sub>S2-2</sub> | 83.94 ± 11.26 | 84.77 ± 9.06 | 85.05 ± 6.15 | 85.44 ± 5.48 | 84.64 ± 4.10 |
| | W<sub>Comb-2</sub> | 85.77 ± 5.40 | 86.34 ± 3.62 | 85.06 ± 4.26 | 87.90 ± 6.43 | 84.56 ± 3.54 |
| Sense of Urgency (APCPS+Mean FixDur) | SoU<sub>S1</sub> | 51.08 ± 10.23 | 53.19 ± 12.06 | 51.24 ± 9.17 | 52.54 ± 9.55 | 49.52 ± 12.26 |
| | SoU<sub>S2</sub> | 61.05 ± 11.14 | 62.43 ± 8.27 | 56.02 ± 9.79 | 61.77 ± 7.98 | 58.94 ± 9.09 |
| | SoU<sub>Comb</sub> | 58.19 ± 7.66 | 61.32 ± 5.91 | 52.91 ± 5.37 | 59.34 ± 5.46 | 58.31 ± 5.95 |
| Mind Wandering (APCPS+EEG) | MW<sub>S1</sub> | 69.80 ± 11.35 | 73.16 ± 8.92 | 74.93 ± 7.38 | 73.02 ± 5.55 | 74.36 ± 6.27 |
| | MW<sub>S2</sub> | 85.40 ± 7.65 | 83.29 ± 9.43 | 81.24 ± 3.76 | 83.91 ± 2.15 | 82.87 ± 6.72 |
| | MW<sub>Comb</sub> | 79.09 ± 4.90 | 77.23 ± 4.30 | 70.97 ± 6.74 | 78.09 ± 4.49 | 76.66 ± 6.79 |

Note that combining EEG and pupillometry to characterize the cognitive states has been previously considered in [1, 2, 29, 86], and a few of them stated that combining these two signals may not lead to any improvement in cognitive state classification accuracy (*e.g.*, see [86]). Since combining APCPS and EEG not only decreases classification accuracy but also potentially increases running time due to the higher number of features, we conclude that there does not seem to be a clear benefit from integrating multiple physiological signals.

### Statistical Performance Analyses

We further performed statistical analyses on the five ML algorithms to determine which of them had the overall highest performance. First, we ran the five ML algorithms for four classification tasks including 3-class workload classification, 2-class workload classification, sense of urgency classification, and mind wandering classification by using the APCPS 19 times each and determined the accuracy arrays obtained from 19 runs. Then, we calculated the (Mean + SD) of the accuracy arrays corresponding to four classification tasks in Table 16 (upper table). After that, we performed full ANOVA model test to investigate the performance variations of five ML algorithms in four classification tasks (the results were presented in Table 16 below table) which showed significant outcomes for 2-class workload classification, 3-class workload classification, and mind wandering classification (p-values *<* .05). Afterwards, we ran a post-hoc analyses by using the paired sampled t-tests performed on the accuracy arrays obtained from five ML algorithms. Table 17a, Table 17b, Table 17c, and Table 17d depict the outcomes of the paired sampled t-tests performed on the accuracy arrays acquired from five ML classifiers corresponding to 3-class workload classification, 2-class workload classification, sense of urgency classification, and mind wandering classification, respectively. In particular, for a given type of signal (the APCPS) and a learning algorithm, we repeated the experiment 19 times (each had the 5-fold and provided one accuracy in the test) to collect 19 classification accuracies. Then, we run the paired sampled t-test and obtained the p-values (at a significance level of .95) to investigate the similarities between different pairs of ML algorithms. In addition, we presented the mean of the accuracy arrays acquired from five ML classifiers in Table 16 along with Table 17 to further determine the best and the worst algorithms. The results indicated that we can differentiate RF from all of the other four algorithms in the case of 3-class workload classification and mind wandering classification. On the other hand, there is no consistency in the p-values related to 2-class classification and sense of urgency classification. In overall, we observed that except for RF (in 3-class workload classification and mind wandering classification), there is no statistically significant difference between the accuracies of the ML algorithms used in this study for classifying different workload and mind wandering levels. Based on Table 16, we can claim that RF is the worst algorithm in achieving different cognitive state classification tasks.

**Table 16.** (Above table) The mean and the standard deviations of the accuracy arrays obtained from five ML classifiers to evaluate the similarities of their effects on (a) W_Comb-1_: 3-class workload classification, (b) W_Comb-2_: 2-class workload classification, (c) SoU_Comb_: sense of urgency classification, and (d) MW_Comb_: mind wandering classification. (Below table) The results of the full ANOVA model test performed on all five ML accuracy arrays reported for (a) W_Comb-1_: 3-class workload classification, (b) W_Comb-2_: 2-class workload classification, (c) SoU_Comb_: sense of urgency classification, and (d) MW_Comb_: mind wandering classification.

| | (Mean ± SD)<br>of k-NN | (Mean ± SD)<br>of NB | (Mean ± SD)<br>of RF | (Mean ± SD)<br>of SVM | (Mean ± SD)<br>of MLP |
| --- | --- | --- | --- | --- | --- |
| $W_{\text{Comb-1}}$ | 62.78 ± 5.47 | **63.35 ± 5.63** | 53.64 ± 4.15 | **63.35 ± 4.43** | 63.25 ± 5.22 |
| $W_{\text{Comb-2}}$ | **90.18 ± 3.42** | 87.90 ± 6.54 | 85.35 ± 5.97 | 88.76 ± 4.14 | 85.71 ± 4.41 |
| $\text{SoU}_{\text{Comb}}$ | 57.04 ± 6.52 | **60.46 ± 9.29** | 54.05 ± 5.95 | 57.04 ± 8.41 | 55.75 ± 7.80 |
| $\text{MW}_{\text{Comb}}$ | **82.08 ± 7.35** | 80.51 ± 6.54 | 71.83 ± 7.61 | 77.24 ± 5.70 | 79.94 ± 6.98 |

|  | f-value | p-value | Partial Eta Squared |
| --- | --- | --- | --- |
| $W_{\text{Comb-1}}$ | 8.18 | $1.13 \times 10^{-5}$ | 0.27 |
| $W_{\text{Comb-2}}$ | 9.90 | $1.10 \times 10^{-6}$ | 0.31 |
| $\text{SoU}_{\text{Comb}}$ | 1.68 | 0.16 | $0.69 \times 10^{-1}$ |
| $\text{MW}_{\text{Comb}}$ | 6.18 | $1.94 \times 10^{-4}$ | 0.22 |

**Table 17.** The results of the paired sampled t-test on the accuracy arrays obtained from five ML classifiers to evaluate the similarities of their efficiencies on (a) W_Comb-1_: 3-class workload classification, (b) W_Comb-2_: 2-class workload classification, (c) SoU_Comb_: sense of urgency classification, and (d) MW_Comb_: mind wandering classification. Here, we marked the cells with red which include p-values less than .05 (statistically significant).

|  | k-NN | NB | RF | SVM |
| --- | --- | --- | --- | --- |
| NB | 0.77 | - | - | - |
| RF | $1.7 \times 10^{-4}$ | $5.3 \times 10^{-4}$ | - | - |
| SVM | 0.74 | 0.99 | $3.3 \times 10^{-5}$ | - |
| MLP | 0.78 | 0.96 | $4.3 \times 10^{-4}$ | 0.96 |

(a) $W_{\text{Comb-1}}$ .
|  | k-NN | NB | RF | SVM |
| --- | --- | --- | --- | --- |
| NB | 0.21 | - | - | - |
| RF | **0.01** | 0.25 | - | - |
| SVM | 0.17 | 0.61 | $9.0 \times 10^{-2}$ | - |
| MLP | $3.6 \times 10^{-3}$ | 0.26 | 0.79 | $9.8 \times 10^{-2}$ |

(b) $W_{\text{Comb-2}}$ .
|  | k-NN | NB | RF | SVM |
| --- | --- | --- | --- | --- |
| NB | 0.21 | - | - | - |
| RF | 0.18 | **0.03** | - | - |
| SVM | 0.99 | 0.21 | 0.16 | - |
| MLP | 0.61 | 0.06 | 0.51 | 0.67 |

(c) $\text{SoU}_{\text{Comb}}$ .
|  | k-NN | NB | RF | SVM |
| --- | --- | --- | --- | --- |
| NB | 0.46 | - | - | - |
| RF | $1.3 \times 10^{-3}$ | $4.8 \times 10^{-3}$ | - | - |
| SVM | $2.6 \times 10^{-2}$ | 0.12 | **0.04** | - |
| MLP | 0.32 | 0.76 | $1.5 \times 10^{-3}$ | 0.14 |

(d) $\text{MW}_{\text{Comb}}$ .

### Comparison the Performance of Pupillometry, Arterial Blood Pressure, and Skin Conductance in Classifying Different Cognitive States

To further investigate the efficiency of the APCPS for different systemic human cognitive state classification tasks, we compared our method with the technique proposed in [98]. This paper presented an ML methodology based on ensemble learning to investigate cognitive load differentiation by using different physiological biomarkers involving heart rate (HR), NN interval, skin temperature, and skin response. The authors introduced two-step cognitive load classification: (1) reconstruction of the discriminative features from the original data and (2) performing ensemble model on the data to accomplish cognitive load classification. In our study, we utilized the same physiological biomarkers used in Borisov *et. al* (other than skin temperature as our dataset does not include this feature) as well as the APCPS, and compared the performance of the APCPS with the HR, the NN interval, and the skin conductance for different classification tasks. Our results, reported in Table 18, indicate that the APCPS outperforms among four different physiological biomarkers in four classification tasks including 2-class workload, 2-class workload, sense of urgency, and mind wandering classification. We further analyzed the effectiveness of combining the two most significant features (APCPS and mean HR) in achieving four classification tasks. Our findings thus show that APCPS alone achieves greater accuracy in assessing multiple cognitive state levels than the combination of the two features.

**Table 18.** Classification accuracies (in %) of APCPS, mean HR, mean NN, mean skin conductance, and the combination of APCPS and mean HR (the best two features) for two-class workload, three-class workload, two-class mind wandering, and two-class sense of urgency classification tasks defined in Table 9 by different ML methods.

| Classification Task | Signal Modality | Feature | k-NN | NB | RF | SVM | MLP |
| --- | --- | --- | --- | --- | --- | --- | --- |
| $W_{\text{Comb-1}}$ | Pupillometry | APCPS | $69.14 \pm 6.06$ | $61.99 \pm 4.58$ | $59.75 \pm 3.39$ | $69.14 \pm 5.39$ | $65.37 \pm 0.58$ |
| | ABP | Mean HR | $34.57 \pm 7.71$ | $31.36 \pm 5.49$ | $38.27 \pm 3.28$ | $35.55 \pm 5.24$ | $33.73 \pm 0.14$ |
| | | Mean NN | $37.28 \pm 2.72$ | $31.85 \pm 4.06$ | $35.31 \pm 5.18$ | $34.81 \pm 3.14$ | $32.54 \pm 0.20$ |
| | Skin Conductance | Mean SC | $27.41 \pm 5.02$ | $27.65 \pm 5.35$ | $30.62 \pm 6.52$ | $31.85 \pm 7.26$ | $33.58 \pm 0.07$ |
| | Pupillometry + ABP | APCPS + Mean HR | $63.70 \pm 5.44$ | $60.96 \pm 6.11$ | $58.68 \pm 6.38$ | $61.81 \pm 7.03$ | $63.70 \pm 6.02$ |
| $W_{\text{Comb-2}}$ | Pupillometry | APCPS | $89.63 \pm 4.57$ | $90.74 \pm 3.77$ | $87.04 \pm 2.92$ | $88.52 \pm 5.24$ | $86.99 \pm 5.33$ |
| | ABP | Mean HR | $48.15 \pm 9.18$ | $44.44 \pm 9.69$ | $53.33 \pm 5.21$ | $50.37 \pm 7.10$ | $50.33 \pm 1.00$ |
| | | Mean NN | $54.07 \pm 8.43$ | $44.81 \pm 4.19$ | $50.37 \pm 8.08$ | $48.52 \pm 6.69$ | $50.00 \pm 1.00$ |
| | Skin Conductance | Mean SC | $46.66 \pm 6.66$ | $43.70 \pm 8.23$ | $42.59 \pm 6.63$ | $45.93 \pm 4.91$ | $50.33 \pm 1.00$ |
| | Pupillometry + ABP | APCPS + Mean HR | $83.15 \pm 4.74$ | $82.41 \pm 4.91$ | $81.77 \pm 4.16$ | $82.15 \pm 5.24$ | $85.30 \pm 2.92$ |
| $MW_{\text{Comb}}$ | Pupillometry | APCPS | $79.94 \pm 4.68$ | $80.25 \pm 5.15$ | $79.99 \pm 8.88$ | $81.66 \pm 7.03$ | $78.88 \pm 3.97$ |
| | ABP | Mean HR | $52.47 \pm 8.43$ | $56.48 \pm 7.17$ | $55.93 \pm 7.16$ | $46.66 \pm 7.03$ | $51.11 \pm 1.84$ |
| | | Mean NN | $52.77 \pm 2.93$ | $51.86 \pm 6.14$ | $48.52 \pm 6.69$ | $50.00 \pm 8.16$ | $50.28 \pm 0.83$ |
| | Skin Conductance | Mean SC | $48.14 \pm 8.92$ | $48.46 \pm 8.70$ | $42.59 \pm 6.99$ | $46.66 \pm 5.67$ | $49.16 \pm 2.50$ |
| | Pupillometry + ABP | APCPS + Mean HR | $76.62 \pm 6.39$ | $73.62 \pm 5.82$ | $74.69 \pm 6.20$ | $74.47 \pm 7.26$ | $74.38 \pm 7.38$ |
| $SoU_{\text{Comb}}$ | Pupillometry | APCPS | $71.30 \pm 8.88$ | $67.27 \pm 7.72$ | $69.93 \pm 4.38$ | $70.22 \pm 9.30$ | $65.61 \pm 5.88$ |
| | ABP | Mean HR | $45.59 \pm 6.00$ | $46.91 \pm 9.02$ | $48.52 \pm 7.72$ | $47.41 \pm 10.97$ | $49.44 \pm 2.08$ |
| | | Mean NN | $49.69 \pm 7.80$ | $50.31 \pm 4.98$ | $48.88 \pm 7.20$ | $48.15 \pm 8.76$ | $50.00 \pm 0.00$ |
| | Skin Conductance | Mean SC | $41.20 \pm 8.22$ | $43.52 \pm 6.28$ | $43.33 \pm 5.44$ | $48.88 \pm 3.85$ | $49.44 \pm 3.69$ |
| | Pupillometry + ABP | APCPS + Mean HR | $68.14 \pm 8.33$ | $61.60 \pm 6.52$ | $67.21 \pm 7.76$ | $66.06 \pm 7.61$ | $85.30 \pm 2.92$ |

### Summary of the Observations

Given the numerical results from the workload, sense of urgency, and mind-wandering classification tasks, we can summarize the main observations as follows:

1. There is no difference in performance among the five machine learning algorithms for a given type of signal and classification task except for RF which had lower accuracies for both 3-class workload classification and mind wandering classification tasks (see Tables 17a, 17d, and 16).
2. APCPS outperforms all other physiological signals in differentiating three cognitive states by a large margin for all six workload, three sense of urgency, and three mind wandering levels regardless of learning algorithms.
3. Combining the APCPS and EEG signals does not really boost the classification performances in workload, sense of urgency, and mind-wandering classification tasks.

It is worth noting that our numerical results from the machine learning models also agreed with our statistical analysis performed on the APCPS signal. As previously mentioned, because of the training effect in session 1 where participants first encountered the primary driving task and secondary tasks, leading to more attention and awareness, one may expect a higher workload in session 1 compared to session 2, specifically during l_w1_ where the participants were first instructed to complete the braking events and dialogue communications (see Figure S7 Fig). This has been observed in our modeling results, for example, by comparing the classification accuracy of the APCPS in task W_S1_ *_−_*_1_ (Table 10a) and in task W_S2_ *_−_*_1_ (Table 10b). As seen, for a given task of classifying between three workload levels, a classifier based on the APCPS signal extracted from session 1 can achieve the highest accuracy of 72.57 ∓ 8.75, while the highest accuracy of the APCPS signal extracted from session 2 is far below which is just 59.13 ∓ 8.29. A similar observation can be made by comparing the classification accuracy of APCPS in task W_S1_ *_−_*_2_ (Table 11a) and in task W_S2_ *_−_*_2_ (Table 11b). Particularly, for a given task of classifying between the highest and lowest workload levels, a classifier based on the APCPS signal extracted from session 1 can achieve the highest accuracy of 93.25 ∓ 3.55 in task W_S1_ *_−_*_2_ which is, again, a bit higher than the highest accuracy of 88.89 ∓ 5.23 on the APCPS signal extracted from session 2 in task W_S2_ *_−_*_2_.

## Discussion

We have explored in great detail the utility of multiple physiological signal modalities based on pupillometry, EEG, fNIRS, skin conductance and respiration for assessing three systemic human cognitive states (cognitive workload, sense of urgency, and mind wandering) by using multiple pairwise statistical tests and state-of-the-art machine learning models. Our statistical analyses results demonstrated that APCPS is the best physiological signal for distinguishing the three systemic cognitive states. We then examined the performance of various machine learning models (k-NN, NB, RF, SVM, and MLP) on the full set of physiological signal types for differentiating different levels of workload, sense or urgency, and mind wandering. Specifically, we performed four cognitive state classification tasks including 3-class workload, 2-class workload, 2-class sense of urgency, and 2-class mind wandering classification which showed that APCPS is the most reliable physiological indicator for achieving high performance in all four classification tasks regardless of which machine learning method was used. In addition to investigating the performance of individual physiological signal modalities in assessing different human cognitive state levels, we also analyzed the efficiency of combining the best two signal types (eye gaze and EEG) for the prediction of different human cognitive states and observed a similar phenomenon: that fusing different physiological signal markers did not improve the classification performance compared to only using eye gaze.

Besides individual cognitive state estimation tasks, we also investigated the association between workload and sense of urgency. Particularly, we analyzed how sequential occurrences of the sense of urgency might impact the overall workload. We considered the APCPS as the most suitable physiological marker (based on its superior performance on cognitive state classification tasks) for exploring the relationship between workload and sense or urgency, and found that consecutive triggering of sense of urgency contributed to overall workload increase. In addition, we examined the effect of possible arousal in sense of urgency prediction. Table 19 includes the p-values and the f-values of Tukey’s HSD multiple pairwise comparison test for different physiological biomarkers obtained from pupillometry, arterial blood pressure, and skin conductance, and compare their efficiency in differentiating sense of urgency levels. The results indicated that the APCPS, taken from pupillometry, is the only biomarker to distinguish the two sense of urgency levels. Physiological biomarkers obtained from skin conductance (mean skin conductance) and arterial blood pressure (mean NN and mean HR) are more associated with individuals’ physiological arousal whereas eye gaze features, taken from pupillometry, are more vulnerable to cognitive state variation such as sense of urgency or attention. Fluctuations in skin conductance and ABP often reflect sympathetic nervous system arousal and are more associated with physiological activity [76]. Sense of urgency may not always stimulate notable changes in biomarkers acquired from skin conductance and ABP given that sense of urgency is considered as a more situational response whereas arousal may remain consistent for longer time periods. On the other hand, pupillometry is specifically sensitive to cognitive variations including cognitive effort and attention which are more related to sense of urgency. Sense of urgency is correlated with how much cognitive resources are being allocated as a response to time pressure or task requirements. Therefore, pupillometry is expected to have better performance in differentiating among multiple levels of sense of urgency occurrences. Considering that our results indicates superior performance of pupillometry and worse performance of skin conductance and ABP in classifying pre-defined sense of urgency levels, this suggests that the pupillometry response is more vulnerable to the cognitive demands of the individuals (which are related to sense of urgency) rather than general physiological arousal. Hence, the superior efficiency of pupillometry over skin conductance and ABP in differentiating these two levels indicates that pupillometry is effectively classifying sense of urgency levels rather than arousal. Pupillometry, in particular PCPS, has the ability to capture various attentional and cognitive variations which makes it an effective physiological biomarker for distinguishing multiple workload, sense of urgency, and mind wandering levels. Although all of these mental states contain an individual’s cognitive effort and attentional fluctuations, they differ significantly across several aspects. First, workload is linked to mental demands imposed by task requirements and tends to increase over an extended period of time. Second, the sense of urgency is associated with an individual’s extended focus due to time pressure and the demand for quick decision-making. Third, mind wandering is defined as a specific type of spontaneous thought unrelated to the task at hand which lies between creative thinking and dreaming. Even though all three cognitive states reflect overlapping processes (*e.g.,* higher levels of mind wandering are often associated with lower perceived workload), they can be differentiated based on their underlying cognitive and attentional mechanisms. Workload is primarily driven by the demands of the task and tends to accumulate gradually over time. Sense of urgency reflects heightened sustained attention due to external time pressure and the need for rapid responses. Mind wandering, in contrast, involves internally generated thoughts that are decoupled from the task and can occur spontaneously, independent of immediate task demands. These distinctions indicate that, while pupillometry captures shared aspects of cognitive effort and attentional fluctuation, it also provides sensitivity to the unique temporal and functional characteristics of each state, enabling it to discriminate between them in different contexts.

**Table 19.** p-values and f-values of Tukey HSD multiple pairwise test for the physiological biomarkers obtained from pupillometry, ABP, and skin conductance for two sense of urgency levels.

| Signal Type | Feature | f-value of $l_{\text{sou}_1} - l_{\text{sou}_2}$ | p-value of $l_{\text{sou}_1} - l_{\text{sou}_2}$ |
| --- | --- | --- | --- |
| Pupillometry | APCPS | 39.31 | $2.66 \times 10^{-9}$ |
| ABP | Mean NN | 0.25 | 0.62 |
|  | Mean HR | 0.22 | 0.64 |
| Skin Conductance | Mean Skin Conductance | 0.02 | 0.87 |

Finally, we performed a statistical comparison of different machine learning algorithms, repeating the experiment multiple times by using the APCPS (each time we performed 5-fold cross validation and obtained one accuracy in the test) for four classification tasks (defined in Model Development and Evaluation section) and ran paired *t*-test within five ML algorithms. We found that except for RF, the remainder of four ML algorithms show roughly equal performance in terms of 3-class workload and 2-class sense of urgency classification accuracy. Moreover, all of the five ML algorithms are comparable in accomplishing 2-class workload and 2-class mind wandering classification. The results indicate that APCPS is able to achieve high prediction performance irrespective of which ML method is used. This result is especially important for ambulatory humans in mixed-initiative teams where certain types of physilogical sensor instrumentations are not possible (*e.g.*, EEG and fNIRS would cause large motion artifacts that might overshadow any changes in the signals that could reflect cognitive state changes). Being able to use eye gaze to track the human workload and sense of urgency as well as mind wandering will allow autonomous systems to better estimate human situational awareness, to adapt their interactions based on the detect levels of human systemic cognitive states, and to utilize opportunities to intervene to address possible degradation of human performance.

### Limitations and Future Work

The outcomes of this study are important for future efforts to assess systemic human cognitive states in real-time considering that human gaze is easy to collect and fast to process compared to other physiological signals. To do so, future work will need to develop methods for real-time assessment and tracking of these human cognitive states, and possibly additional states such as interference and distraction which have been shown to impact human task performance as well. It is also worth pointing out that the dataset utilized in this study was collected from participants who were mostly undergraduate and graduate students with an average age of just 20. It would be important to verify that the results obtained based on this dataset also hold for a broader set of human participants for whom we expect to see even greater variations in the various cognitive state levels. And we need to investigate a wider set of systemic cognitive states such as distraction (*e.g.*, [88] or interference (*e.g.*, [89]) to further explore the utility of pupillometry. Finally, while the characteristics of current task setting (ongoing manual actions based on visual perceptions, intermittent spoken dialogue interactions) was an instance of a fairly large class of tasks, it would be interesting to consider other tasks with different characteristics to see whether the results would transfer.

## Conclusions

The aim of this comprehensive analysis and modeling effort was to investigate the utility of various physiological signal types such as pupillometry, EEG, fNIRS, skin conductance, and respiration for assessing different human systemic cognitive states like workload, sense of urgency, and mind wandering in a multimodal driving simulation environment. Our statistical analyses and classification results indicate that the “percent change in pupil size” is the single most reliable physiological marker for predicting different levels of workload, sense of urgency and mind wandering. We also explored the change in workload relative to changes in sense of urgency by utilizing the PCPS and determined that frequent and consecutive instances of higher sense of urgency are likely to increase the overall workload. Based on the significant differences in levels of systemic cognitive states, we examined the performance of five different state-of-the-art machine learning models to determine which would be best in predicting those levels and found that except for one (“random forest”), all other four models (k-neareast neighbor, naive Bayes, support vector machines, and muli-layer perceptron) are equally good at classifying the various levels of three systemic cognitive states, critically in data from novel subjects on which these models were not trained, thus showing their potential for state classification in general. While the results were obtained using data from only one task, we believe that they have the potential to generalize to other tasks and thus for improving human performance in safety-critical applications as well as in human-machine teams.

## Supporting information

**S1 Fig. The relationship between Different Cognitive State Pairs.** An illustration of the hypothesized relationships among pairs of three systemic cognitive states: workload, sense of urgency, and mind wandering.

**S2 Fig. An illustration of a braking event.** An example of a braking event: (a) a participant drives on the right lane of a straight road, (b) a car appears in front of the participant’s car, and (c) the front car immediately decelerates, then moves far away.

**S3 Fig. Evaluation of three cognitive workload levels.** By selecting different combinations of braking and dialogue events, three different cognitive workload levels were generated.

**S4 Fig. Evaluation of two sense of urgency levels.** By combining different urgency moments of braking and dialogue events, two sense of urgency levels were generated.

**S5 Fig. The cross correlation depiction between the EEG signal before and after applying the Kalman Smoother**. The investigation of impact of the Kalman Smoother on the EEG signals.

**S6 Fig. The change in the mean APCPS related to different cognitive workload levels.** Boxplot of the mean APCPS values corresponding to different cognitive workload levels including both sessions from all of the subjects.

**S7 Fig. The change in the mean APCPS related to different cognitive workload levels for two sessions.** Boxplot of the mean APCPS values corresponding to different sessions of three cognitive workload levels.

**S8 Fig. The change in the mean APCPS related to different sense of urgency levels.** Boxplot of the mean APCPS corresponding to two sense of urgency levels including both sessions from all of the subjects. l_sou1_: Lower sense of urgency, l_sou2_: Higher sense of urgency.

**S9 Fig. The change in the mean APCPS related to different mind wandering levels.** Boxplot of the mean APCPS corresponding to two mind wandering levels including both sessions from all of the subjects. l_mw1_: Lower mind wandering, l_mw2_: Higher mind wandering.

**S10 Fig. Illustration of the change in PCPS with different events.** Illustration of the change in PCPS with different events including braking and dialogue. The results include two sessions obtained from two different subjects. B and D represent the onset of the braking event and the onset of the dialogue event, respectively.

## Acknowledgments

This project was in part supported by AFOSR grant #FA9550-18-1-0465.

